# Organization of Peptidoglycan Synthesis in Nodes and Separate Rings at Different Stages of Cell Division of *Streptococcus pneumoniae*

**DOI:** 10.1101/2020.09.27.315481

**Authors:** Amilcar J. Perez, Michael J. Boersma, Kevin E. Bruce, Melissa M. Lamanna, Sidney L. Shaw, Ho-Ching T. Tsui, Atsushi Taguchi, Erin E. Carlson, Michael S. VanNieuwenhze, Malcolm E. Winkler

**Affiliations:** Department of Biology, Indiana University Bloomington, Bloomington, IN 47405 USA; Department of Microbiology, Harvard Medical School, Boston, MA 02115 USA; Department of Chemistry, University of Minnesota, Minneapolis, MN 55455 USA; Department of Chemistry, Indiana University Bloomington, Bloomington, IN 47405 USA

**Keywords:** septal and peripheral peptidoglycan synthesis, penicillin-binding proteins (PBPs), FtsX, fluorescent D-amino acids (FDAAs), high-resolution 3D-SIM

## Abstract

Bacterial peptidoglycan (PG) synthesis requires strict spatial and temporal organization to reproduce specific cell shapes. In the ovoid-shaped, pathogenic bacterium *Streptococcus pneumoniae* (*Spn*), septal and peripheral (sidewall-like) PG synthesis occur simultaneously at midcell. To uncover the organization of proteins and activities that carry out these two modes of PG synthesis, we examined *Spn* cells vertically oriented onto their poles to image the division plane at the high lateral resolution of 3D-SIM (structured-illumination microscopy). Using fluorescent D-amino acid (FDAA) probes, we show that areas of new transpeptidase (TP) activity catalyzed by penicillin-binding proteins (PBPs) separate into a pair of concentric rings early in division, representing peripheral PG (pPG) synthesis (outer ring) and the leading-edge (inner ring) of septal PG (sPG) synthesis. Fluorescently tagged PBP2x or FtsZ locate primarily to the inner FDAA-marked ring, whereas PBP2b and FtsX remain in the outer ring, suggesting roles in sPG or pPG synthesis, respectively. Short pulses of FDAA labeling revealed an arrangement of separate regularly spaced “nodes” of TP activity around the division site of predivisional cells. Control experiments in wild-type and mutant strains support the interpretation of nodal spacing of TP activity, and statistical analysis confirmed that the number of nodes correlates with different ring diameters. This nodal pattern of FDAA labeling is conserved in other ovoid-shaped species. Tagged PBP2x, PBP2b, and FtsX proteins also exhibited nodal patterns with spacing comparable to that of FDAA labeling. Together, these results reveal a highly ordered PG synthesis apparatus in ovococcal bacteria at different stages of division.

**SIGNIFICANCE:** The spatial organization of PBPs and their TP activity at division septa is not well understood. In some bacteria, TP activity and PBP localization seem to be nodal (also called punctate), whereas in other bacteria, discrete foci of PBP activity are infrequently or not observed. Here we report two basic properties of the organization of PBPs and TP activity in the ovoid-shaped bacterium *Spn*. First, there is distinct spatial separation of the sPG machine, including FtsZ, from the pPG synthesis machine at the midcell of dividing *Spn* cells. Second, in predivisional cells, PBPs and TP activity are organized heterogeneously into regularly spaced nodes, whose number and dynamic distribution are likely driven by the PG synthesis of PBP:SEDS complexes.

## INTRODUCTION

*Streptococcus pneumoniae* (pneumococcus; *Spn*) is a Gram-positive, commensal bacterium of humans and a major opportunistic pathogen that causes life-threatening illnesses, including pneumonia, bacteremia, and meningitis (1, 2). *Spn* is a prolate ellipsoid-shaped bacterium, referred to as an ovococcus (3). Its distinct ovoid shape is maintained by a thick peptidoglycan (PG) wall that surrounds the entire cell (4). PG consists of glycan sugar chains of alternating units of β-(1,4) linked N-acetylglucosamine (GlcNAc) and N-acetylmuramic acid (MurNAc). Glycan chains are crosslinked together by PG peptides attached to MurNAc, thereby forming a mesh-like network (5, 6). As in most eubacteria, *Spn* PG synthesis is essential for normal growth and cell division and determines normal cell shape and chaining, which impact colonization and virulence (4, 7, 8). Besides protecting cells from turgor stress, *Spn* PG serves as a scaffold for surface-attached virulence factors, including capsule, sortase-attached proteins, and wall teichoic acids (4).

In *Spn*, PG synthesis is organized initially by an FtsZ ring at the midcell equator perpendicular to the long axis of newly divided cells (Fig. S1). PG synthesis is carried out by distinct septal PG (sPG) and peripheral PG (pPG) modes (9–11). sPG synthesis produces the cross wall that separates daughter cells (12–14), whereas concurrent pPG synthesis elongates daughter cells from midcell to form ovoid-shaped cells (Fig. S1) (11–14). pPG synthesis in *Spn* functionally resembles preseptal PG elongation that occurs briefly at the beginning of division of rod-shaped bacteria (15, 16). However, pPG elongation in *Spn* cells remains confined to the midcell region, in contrast to lateral PG elongation organized by MreB-containing Rod complexes moving over the body of rod-shaped cells (5).

The location of these two modes of PG synthesis within the same midcell region requires significant coordination and organization. Previously, we used dual-protein 2D-immunofluorescence microscopy (IFM) and high-resolution 3D-SIM to determine the relative localization patterns of pneumococcal division and PG synthesis proteins at different stages of division (10–12). Stages of cell division were assigned retrospectively based on static “snap-shot” images of cells from non-synchronized exponential cultures. These studies showed that in newly divided, predivisional (stage 1) daughter cells, division and PG synthesis proteins locate to an FtsZ-organized, midcell equatorial ring (Fig. S1). As nearly simultaneous sPG and pPG synthesis begin (stage 2) (17), the equatorial ring begins to constrict and becomes the new septal ring. Some FtsZ and associated proteins EzrA and FtsA move outward continuously with MapZ from the septal ring toward the positions of the future equatorial rings in daughter cells (13). As constriction and midcell elongation continue, FtsZ amount decreases at the midcell of middle- to late-divisional cells and begins to accumulate at the equators of daughter cells. Meanwhile, PG synthesis proteins remain at the midcell septal ring.

At the resolution of standard 2D-epifluorescence microscopy (2D-EFM), essential class B bPBP2x transpeptidase (TP), which interacts with the FtsW SEDS glycosyltransferase (GT) to carry out sPG synthesis (13, 18), locates in an apparent inner ring within an outer ring containing the class B bPBP2b TP (11), which interacts with the RodA SEDS GT to carry out pPG synthesis (Fig. S1) (19, 20). Several other proteins implicated in pPG synthesis locate to this apparent outer ring, including MreC that organizes and may activate bPBP2b (21), class A aPBP1a that has both GT and TP activities (12), MltG endo-lytic transglycosylase that cleaves glycan chains (21), and StkP serine/threonine kinase that mediates cell division (11, 12). Near the end of cell division (stage 4), remaining PG synthesis proteins that have not moved to developing equatorial rings converge to a small spot between the daughter cells (Fig. S1) (11), before finally moving to the equatorial rings of the daughter cells (13).

Additional evidence for physical separation of the sPG and pPG synthesis machines midway through cell division was obtained by staining with fluorescent vancomycin (FL-Vanco), which labels PG pentapeptides in areas of PG synthesis (22), with fluorescent D-amino acids (FDAAs), which label PG in regions of active PBP TP activity (23, 24),and with an activity-based β-lactone fluorescent probe (7FL) specific for bPBP2x (25, 26). FL-Vanco labeling combined with IFM of mid-divisional cells indicated localization of bPBP2x inside of regions of nascent PG synthesis that were surrounded by bPBP2b and other pPG synthesis proteins (11). Long pulses of FDAAs for 5 min followed by 3D-SIM showed labeling of a midcell ring surrounding a prominent central dot of FDAA labeling, resembling a “Saturn-like” pattern, in mid-divisional cells (11). Specific inhibition with the β-lactam antibiotic methicillin and protein depletion experiments indicated that the central dot of septal TP activity is attributable to bPBP2x (11). Likewise, these central dots of FDAA labeling disappeared in rings of non-constricted septa in cells depleted for the GpsB regulatory protein (10). Last, spatial separation of bPBP2x was confirmed with a bPBP2x-specific, activity-based β-lactone fluorescent probe (7FL, Lac(L)-Phe-FL), whose labeling recapitulated a “Saturn-like” labeling pattern, indicating that although most active bPBP2x moves toward the center of the septum, some still remains in the outer peripheral ring (25, 26).

Limitations of these previous fluorescent-protein and -probe studies are the low resolution (≈250 nm) of conventional 2D-EFM and of rotated 3D-SIM images (27). Due to their ovoid shape, *Spn* cells predominantly lie sideways along their long axis during imaging on a flat surface. Imaging of septal rings and associated structures in horizontal cells by 3D-SIM is limited by the Z-axial resolution of 250-300 nm of the system, which is much lower than the 100 nm resolution in the XY lateral plane (27). This axial resolution limitation remains when reconstructed image volumes of horizontally oriented cells are rotated by 90° to visualize septal rings as circles. To overcome this obstacle, we devised a simple method to reliably orient a small, but sufficient, number of *Spn* cells vertically onto their poles (i.e., “on their heads”) for 3D-SIM imaging, thereby enabling high lateral resolution of division rings. Using this approach, we were able to detect FDAA-labeled intermediate concentric rings of sPG synthesis TP activity in strains expressing fluorescent PG synthesis proteins. This approach located the proteins to the outer peripheral or inner septal rings and showed that these proteins are organized as nodal structures distributed regularly around these rings in early divisional cells. We further found that short pulses of FDAA labeling also revealed nodal distributions of TP activity that were disrupted in division mutants. Together, these results reveal a highly uniform organization of coordinated PG synthesis and distinct separation of the sPG and pPG synthesis machines at the midcell of dividing ovococcal bacteria.

## RESULTS

### “Snap-shots” of vertically oriented *Spn* cells labeled with FDAAs reveal constriction of concentric septal and peripheral rings

To circumvent the lower resolution during rotation of 3D-SIM images, we devised a simple method to capture a sufficient number of vertically oriented, fixed *Spn* cells to image division planes at high ≈100 nm × ≈100 nm XY lateral resolution. Wild-type (WT) D39 ∆*cps* cells were first labeled with a blue FDAA (HADA) for several generations and then with a red FDAA (TADA) for 2.5 min to label sites of new PG synthesis (Fig. 1A and 1B). Cells were fixed with formaldehyde and resuspended in a small amount of hardening anti-fade solution pipetted onto small, round glass coverslips onto which slides were placed. The anti-fade suspension medium minimized photobleaching, and its viscosity and the large number of cells in samples trapped some cells in a vertical orientation. This procedure led to a reliable, but small (≤1%), percentage of cells suspended on their poles, which could be spotted as circles (Fig. 1A and 1C), among many horizontally oriented cells (Fig. 1B) in each field.

**Fig. 1.**
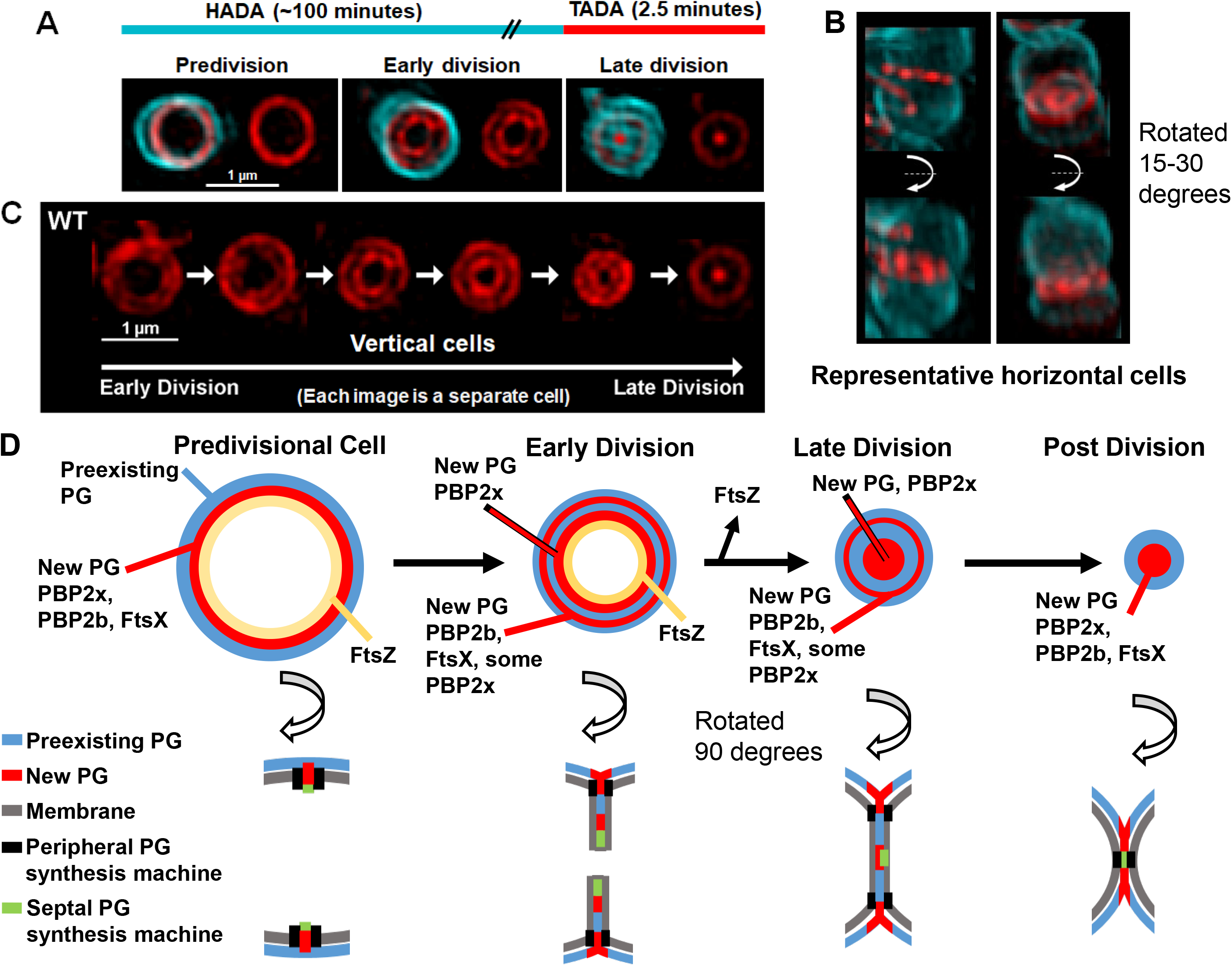
3D-SIM of vertically oriented *Spn* cells labeled with FDAAs reveals spatially distinct, concentric midcell ring intermediates of TP activity at different stages of division. (*A*) Schematic of labeling procedure (*top*). WT cells (IU1945) were labeled with 125 μM HADA for several generations (old cell wall; cyan), washed, and labeled with 125 μM TADA for 2.5 min (new cell wall from ≈7% of a generation; red). Cells were fixed and prepared for vertical cell imaging by 3D-SIM (see *SI Appendix, Experimental Procedures*). Representative images are shown of rings from pre-, early, and late-division stages estimated by the diameters of outer TADA rings. Left images, HADA and TADA channels overlaid; right images TADA channel only. (*B*) Representative horizontally oriented cells from the same field as (*A)* locating the concentric rings at constricting midcell regions. *(C)* Montage of manually sorted *Spn* cells in different stages of cell division. Only the TADA is shown, and images are representative of >50 cells from more than three independent biological replicates. *(D)* Diagrammatic summary based on data in *Results* of the organization of sPG and pPG synthesis in *Spn*, including concentric rings of new PG synthesis, membrane invagination, and proteins located in this study. Curved arrows indicate the image rotations indicated.

Vertically oriented *Spn* cells were captured at different stages of the division cycle (Fig. 1A and 1C). In predivisional cells, red TADA labeling appears as a single ring around the division site (diagrammed in Fig. 1D). Remarkably, as division precedes, TADA labeling is detected in a pair of intermediate concentric rings, which are designated as the outer (peripheral) and inner (septal) rings. The inner ring represents the TP activity at the leading-edge of the closing septum annulus that eventually converges down to a dot, surrounded by the outer ring to form a “Saturn-like” pattern reported before, where the dot results from bPBP2x TP activity (11). In very late divisional cells, all remaining TADA labeling at midcell converges to a single dot at the division site, while new TADA rings appear at the equators of daughter cells. The regions of TP activity, including their relationship to membrane invagination, are summarized in Figure 1D. We conclude that PG synthesis occurs at two distinct, separate locations at the midcell of dividing *Spn* cells.

### bPBP2x and FtsZ primarily locate to the constricting inner septal ring, whereas bPBP2b and FtsX locate to the outer peripheral ring

We reasoned that the spatial separation observed in vertically oriented *Spn* cells could be used to assign PG synthesis proteins to the sPG and/or pPG synthesis machine(s). We labeled strains expressing isfGFP-bPBP2x or sfGFP-bPBP2b with TADA (125 μM) for a moderate time (2.5 min), and then observed TADA and GFP fluorescence in vertically oriented cells (Fig. 2 and S2). isfGFP-bPBP2x or sfGFP-bPBP2b was expressed from its native chromosomal locus, and cells expressing these constructs exhibited minimal discernible growth or cell morphology defects (Fig. S3A and S3B) (13). Western blotting with native antibody showed a single species of each fusion protein (Fig. S3C). Nevertheless, isfGFP-bPBP2x cellular amount is moderately increased (169%) compared to unlabeled WT bPBP2x, whereas sfGFP-bPBP2b is underproduced to only 12-15% of the WT amount, without causing ostensible defects in cell growth and shape in BHI broth (Fig. S3). We characterized two other strains that express fusion constructs without causing defects in growth and morphology. In one strain, iHT-bPBP2x is expressed at ≈164% or ≈88% of WT bPBP2x level when grown in BHI broth or C+Y medium, respectively (Fig. S2C, S2E, and S3 A-C). In the second iHT-bPBP2b//PZn-iHT-bPBP2b merodiploid strain grown in BHI broth containing 0.3 mM Zn^2+^/0.03 mM Mn^2+^, iHT-bPBP2b is expressed at ≈WT bPBP2b level (≈107%; Fig. S2D and S3C).

**Fig. 2.**
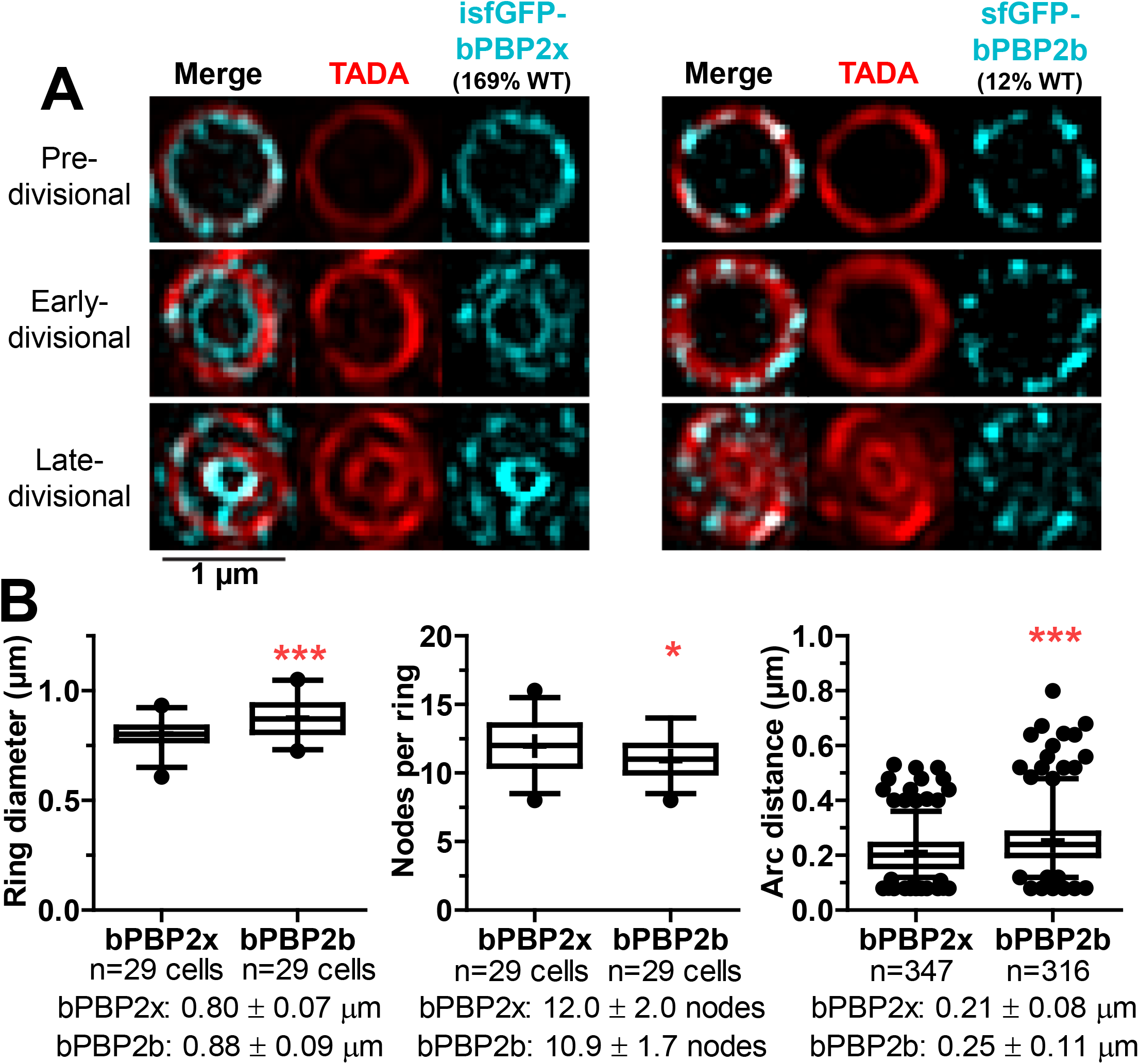
bPBP2x or bPBP2b locate to the leading edge of the septal ring annulus or to the outer peripheral ring, respectively, at midcell in vertically oriented *Spn* cells. Cells expressing isfGFP-bPBP2x (IU11157) or sfGFP-bPBP2b (IU9965) were labeled with 125 μM TADA for 2.5 min, fixed, and prepared for vertical cell imaging by 3D-SIM (see *SI Appendix, Experimental Procedures*). (*A*) Representative images of cells (6 to 29) at different stages of division. isfGFP-bPBP2x and sfGFP-bPBP2b, pseudo-colored cyan; TADA labeling, pseudo-colored red. Brightness and contrast were manually adjusted to show signals associated with division rings and to reduce background lacking TADA labeling. Progression of cell division was indicated by the separation of the inner (septal) and outer (peripheral) rings. Cellular amounts of fluorescent-proteins produced relative to untagged WT protein in a control strain (see Fig. S3C) are in parentheses. (*B*) Quantitation of nodal distributions of isfGFP-bPBP2x and sfGFP-bPBP2b (but not TADA) in predivisional *Spn* cells with single, overlapping rings of TADA and GFP labeling. Distributions of ring diameters (left), nodes per ring (middle), and arc distances (right) were determined as described in *SI Appendix, Experimental Procedures.* Graphs show median (interquartile range; whiskers, 5^th^-95^th^ percentile; +, mean). Means ± SD from two independent biological replicates are shown below the graphs and compiled in Table S3. Differences between means were compared by a two-tailed *t* test (GraphPad Prism). *, *P*<0.05 and ***, *P*<0.001.

In predivisional cells grown in BHI broth, TADA treatment for 2.5 min labels equatorial rings fairly uniformly, whereas sfGFP-bPBP2x and sfGFP-bPBP2b appear as nodes distributed regularly around the circumference of cells (Fig. 2A and, S2A, row 1). Independent labeling of WT bPBP2x with the 7FL β-lactone probe specific for bPBP2x corroborated a nodal organization in predivisional equatorial rings of daughter cells (Fig. S2F). As division progresses, isfGFP-bPBP2x moves inward resulting in formation of the inner septal ring (Fig. 2A and S2A, row 2), with some bPBP2x remaining in the outer ring, as was also observed for labeling with the 7FL (Fig. S2F) (25, 26). Displacement between the single TADA ring and the inner isfGFP-bPBP2x ring (Fig. 2A and S2A, row 2) likely reflects the continued movement inward of isfGFP-bPBP2x during the time required to wash away unincorporated TADA and fix cells. As the TADA-labeled inner septal ring becomes more prominent, isfGFP-bPBP2x condenses into a dense, concentric ring largely lacking detectable nodes at this resolution, with some isfGFP-bPBP2x remaining in the outer ring (Fig. 2A and S2A, row 3).

A similar pattern of localization was observed for slightly overexpressed or underexpressed iHT-bPBP2x in cells grown in BHI broth or C+Y medium, respectively (Fig. S2C and S2E), including residual iHT-bPBP2x remaining in the outer ring. In iHT-protein fusion experiments, the blue FDAA HADA replaced TADA due to spectral overlap of the red HT-JF549 ligand with TADA. Nodes of iHT-bPBP2x (Fig. S2C and S2E) were often less distinct and regular than those of intrinsically fluorescent isfGFP-bPBP2x (Fig. 2A and S2A), likely because of non-optimal labeling with the extrinsic HT substrate.

By contrast, as isfGFP-bPBP2x progresses to the inner septal ring during division (Fig. 2A and S2A, row 3), sfGFP-bPBP2b expressed at a low level compared to WT bPBP2b remains confined to the outer peripheral ring, with little, if any sfGFP-bPBP2b detected in the inner ring marked by TADA (Fig. 2A and S2B, row 3). Likewise, iHT-bPBP2b expressed near the WT bPBP2b level remains as nodes in the outer peripheral ring of pre- and early divisional cells (Fig. S2D, row 1), but is not detected in the constricting inner septal ring (Fig. S2D, row 2). Again, extrinsically labeled iHT-bPBP2b nodes vary more than those of intrinsically fluorescent sfGFP-bPBP2b. Together, these results identify intermediate states of spatial separation of the bPBP2x-containing sPG and bPBP2b-containing pPG synthesis machines during *Spn* division (see Fig. 1D). The nodal localization of bPBP2x and bPBP2b in the midcell rings of predivisional cells is taken up below in the context of FDAA labeling patterns.

We extended this approach to a protein that had not been localized before. FtsX is a polytopic membrane protein that forms a complex with the cytoplasmic FtsE ATPase and the extracellular PcsB PG hydrolase in *Spn* (28–31). The FtsEX:PcsB complex is essential for *Spn* growth due to its role in hydrolytic remodeling during PG synthesis (28, 32, 33). Depletion of FtsEX:PcsB results in the formation of chains of spheroid cells (31, 33), characteristic of defective pPG synthesis (11, 28). To test this idea, we localized an FtsX’-isfGFP-FtsX’ sandwich fusion in vertically oriented *Spn* cells (Fig. S2G). This FtsX sandwich fusion is expressed from the normal chromosomal locus and does not cause cell defects, even though about 60% seemed to be proteolytically cleaved once in the extracellular ECL1 domain near the fusion point (Fig. S3C). Throughout the cell cycle, FtsX-isfGFP-FtsX’ appears as nearly evenly spaced nodes around the outer peripheral ring demarked by TADA labeling and was not detected in the inner septal ring later in division (Fig. S2G). Finally, we determined that FtsZ-sfGFP tracks with TP activity at the leading-edge of the closing septum annulus (Fig. S2H). We conclude that PG remodeling by the FtsEX:PcsB complex is likely confined to pPG synthesis in *Spn*, and that FtsEX and pPG synthesis is not associated with an FtsZ ring during much of the *Spn* cell cycle (see Fig. 1D). In addition, this example illustrates the utility of using FDAA labeling as a fiducial marker to distinguish proteins involved in sPG and/or pPG synthesis.

### New PG synthesis is organized as a series of regularly spaced nodes around the midcell of predivisional *Spn* cells

At a labeling time of 2.5 min with 125 μM TADA, we observed apparent nodal variation of TADA intensity in the outer and inner rings of dividing *Spn* cells (Fig. 1 and 2). As noted above, we also observed nodal positioning of bPBP2x, bPBP2b, and FtsX in these rings (Fig. 2 and S2). To study these patterns further, we labeled WT *Spn* cells with a lower concentration of TADA (45 μM) for a very short pulse (17 s) before imaging vertically oriented cells (Fig. 3). To maximize the spacing between nodes, we confined our analysis of TADA labeling patterns to predivisional cells that have the largest diameters by inspection and that were relatively plentiful in fields of non-synchronized cultures. Very short pulses resolved TADA labeling into a series of regularly spaced nodes distributed around the equators of predivisional *Spn* cells (Fig. 3A).

**Fig. 3.**
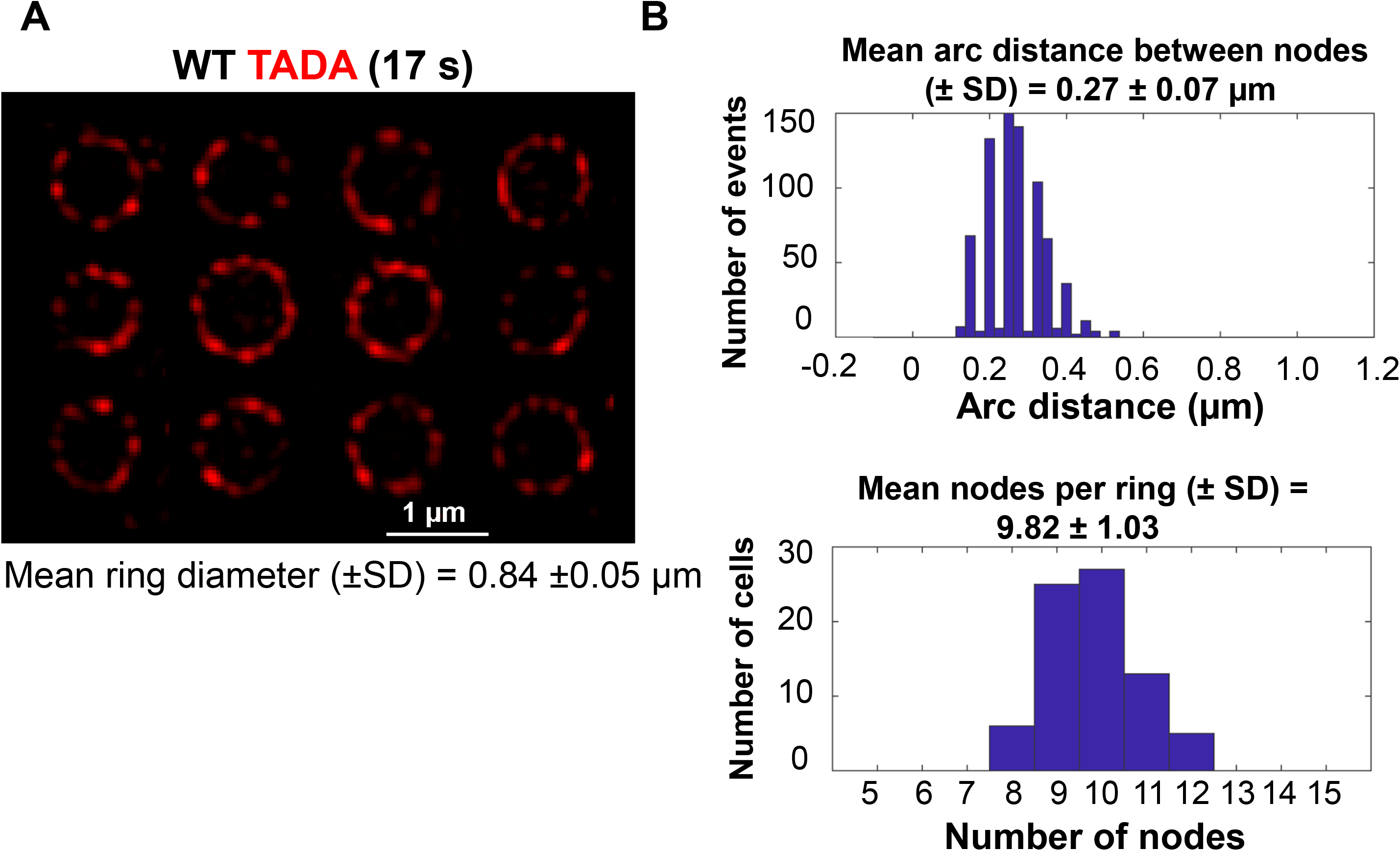
PBP TP activity is organized into regular nodes at the midcell of predivisional *Spn* cells. WT cells (IU1945) were incubated for 17 s with a low concentration of TADA (45.5 μM), fixed, and prepared for vertical cell imaging by 3D-SIM (see *SI Appendix, Experimental Procedures*). (*A*) Representative images of 12 predivisional *Spn* cells pulsed-labeled with TADA. Mean ring diameter ± SD of >60 cells from >3 independent biological replicates was determined as described in *SI Appendix, Experimental Procedures*. (*B*) Distributions of arc distances (*top*) and number of nodes per ring (*bottom*) determined for the data set of cells in (*A*) and compiled in Table S3.

To quantify the pattern of these nodes, we developed a custom vertical image analysis graphical user interface (VIMA-GUI) using MATLAB (see *Experimental Procedures*). After manually designating individual nodes in each division ring, the program accurately determines the location of the ring, the diameter of the cell, the number of FDAA nodes, and the arc distance between adjacent nodes. TADA pulse-labeled and fixed WT ∆*cps* D39 cells have an average diameter of 0.84 μm ± 0.05 (SD), indicating that the selected cells are at the same predivisional stage (Fig. 3A; Table S3). WT ∆*cps* D39 cells contain an average of 9.8 ± 1.0 (SD) individual TADA-labeled nodes, with an average arc distance between nodes of 0.27 μm ± 0.07 (SD) (Fig. 3B; Table S3). The consistency of these measurements strongly supports the notion of an ordered placement of TP activity sites early in *Spn* division.

### The pattern of FDAA-labeled nodes is not caused by image processing

To confirm that the regular nodal TADA-labeling pattern was not caused by the processing steps required for rendering structured illumination images, we examined the raw data and data-process parameters for TADA-pulse-labeled WT ∆*cps* D39 cells (Fig. 1 and 3). Nodal labeling was still observed at the equators of pulse-labeled predivisional *Spn* cells using conventional, low-resolution (≈250 nm) widefield 2D-EFM without image deconvolution (arrows, Fig. 4A). Deconvolution improved the clarity of the nodes in these low-resolution widefield images, but the images remained blurry compared to 3D-SIM images (Fig. 3A). To examine the influence of the major user-defined data smoothing function in SIM reconstruction, we processed 3D-SIM data by adjusting the Wiener filter setting stepwise between 0.001-0.02 (34). Higher filter settings (above 0.01) resulted in clear “honeycomb” patterning effects on the sample and in background regions. We found no discernable patterning or grouping (e.g., over-separation of intensity into nodes) of signal with settings below 0.006 (Fig. 4B, row 2) and set the Wiener filter at 0.001 for the data presented in this work (Fig. 4B, row 1). Finally, we asked if the level of fluorescence signal in the pulse-labeled cells was potentially contributing to an artificial separation of signal into nodes. To this end, we determined the maximal pixel peak intensities in lines drawn through nodes in cells labeled for 17s (Fig. 3) and through the nearly contiguous labeling rings of cells labeled for 2.5 min (Fig. 1). Background offsets were determined from regions lacking cells for the two labeling conditions, and these backgrounds were subtracted from the mean maximal pixel intensities. The mean maximal pixel intensity of nodes (425 A. U. ± 105) was similar to that of nearly contiguous rings (461 A. U. ± 169), indicating that the SIM processing can produce nearly contiguous rings and separate nodes at about the same intensity level. This equivalence argues that the observed nodes are not simply being created by grouping together regions of low fluorescence signal. We conclude that the 3D-SIM processing parameters used in these analyses did not create the TADA nodal patterns observed in vertical *Spn* cells.

**Fig. 4.**
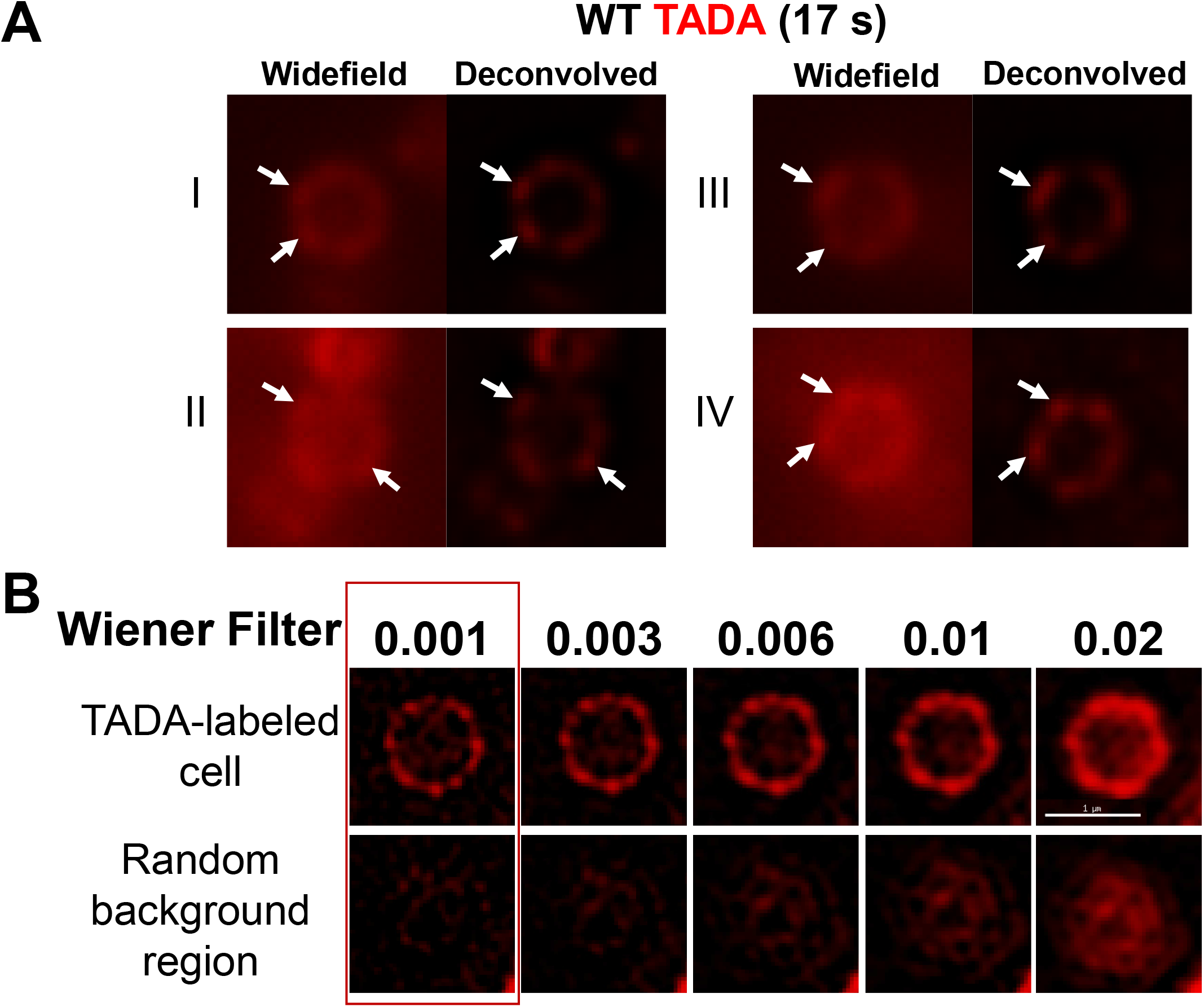
TADA-labeled nodes at the midcell of vertically oriented predivisional *Spn* cells are observed by widefield microscopy and at different 3D-SIM filter settings. WT cells (IU1945) were labeled with 45.5 μM TADA for 17 s, fixed, and prepared for vertical cell imaging as described in *SI Appendix, Experimental Procedures*. (*A*) Representative widefield-microscopy images of four separate cells, before and after deconvolution. Arrows point to nodes of TADA labeling. (*B*) 3D-SIM images processed with different Wiener filter settings of the midcell of a single TADA-labeled vertical cell (top row) or a background region lacking a cell (bottom row). Red box, 0.001 Wiener filter setting used throughout this paper for 3D-SIM imaging. Scale bar = 1 μm.

In support of this conclusion, the nodal pattern of TADA labeling becomes irregular in mutants defective in FtsZ ring assembly or in the regulation of PG synthesis (next section). This irregularity is not consistent with an image reconstruction artifact. In one experiment, FtsZ was expressed from a Zn2+-inducible promoter in a ∆*ftsZ*//PZn-*ftsZ*^+^ merodiploid strain grown in BHI broth containing sufficient Zn^2+^ to allow growth comparable to the WT strain. The merodiploid strain was shifted to BHI broth lacking Zn^+2^ for 2.5 h, which resulted in enlarged spheroid cells with increased (≈1.4-fold) diameters (Fig. 5; Table S3). The FtsZ-depleted and WT FtsZ^+^ strain were pulse-labeled (17 s) with TADA, and vertically oriented cells were imaged by 3D-SIM. WT cells show a regular nodal pattern in ≈94% of vertical cells, whereas the FtsZ-depleted cells with larger diameters show a regular nodal pattern in only ≈40% of cells (Fig. 5A). The remaining ≈60% of FtsZ-depleted cells show irregular nodal arrangements, often with large gaps between nodes (arrow, Fig. 5A). “Regular” and “irregular” nodal patterns were initially distinguished by visual heuristic criteria, where “regular” refers to nodal patterns that appear by eye to be relatively evenly distributed, whereas “irregular” refers nodal distributions that contain one or more large gaps with spacing estimated to be at least twice that in the regular nodal pattern. Subsequent measurements of arc lengths generally matched the initial visual criteria. FtsZ-depleted cells with regular patterns contained more TADA-labeled nodes than WT or FtsZ-depleted cells with irregular patterns (Fig. 5B, middle; Table S3). The mean distance between nodes was greater for FtsZ-depleted than WT cells, with the variability of gap distances in FtsZ-depleted cells with irregular patterns reflected by a high standard deviation (Fig. 5B, right; Table S3). Taken together, this evidence strongly argues that the FDAA nodes are not an artifact of 3D-SIM imaging and that FtsZ is required to maintain the WT arrangement of FDAA nodes in predivisional *Spn* cells.

**Fig. 5.**
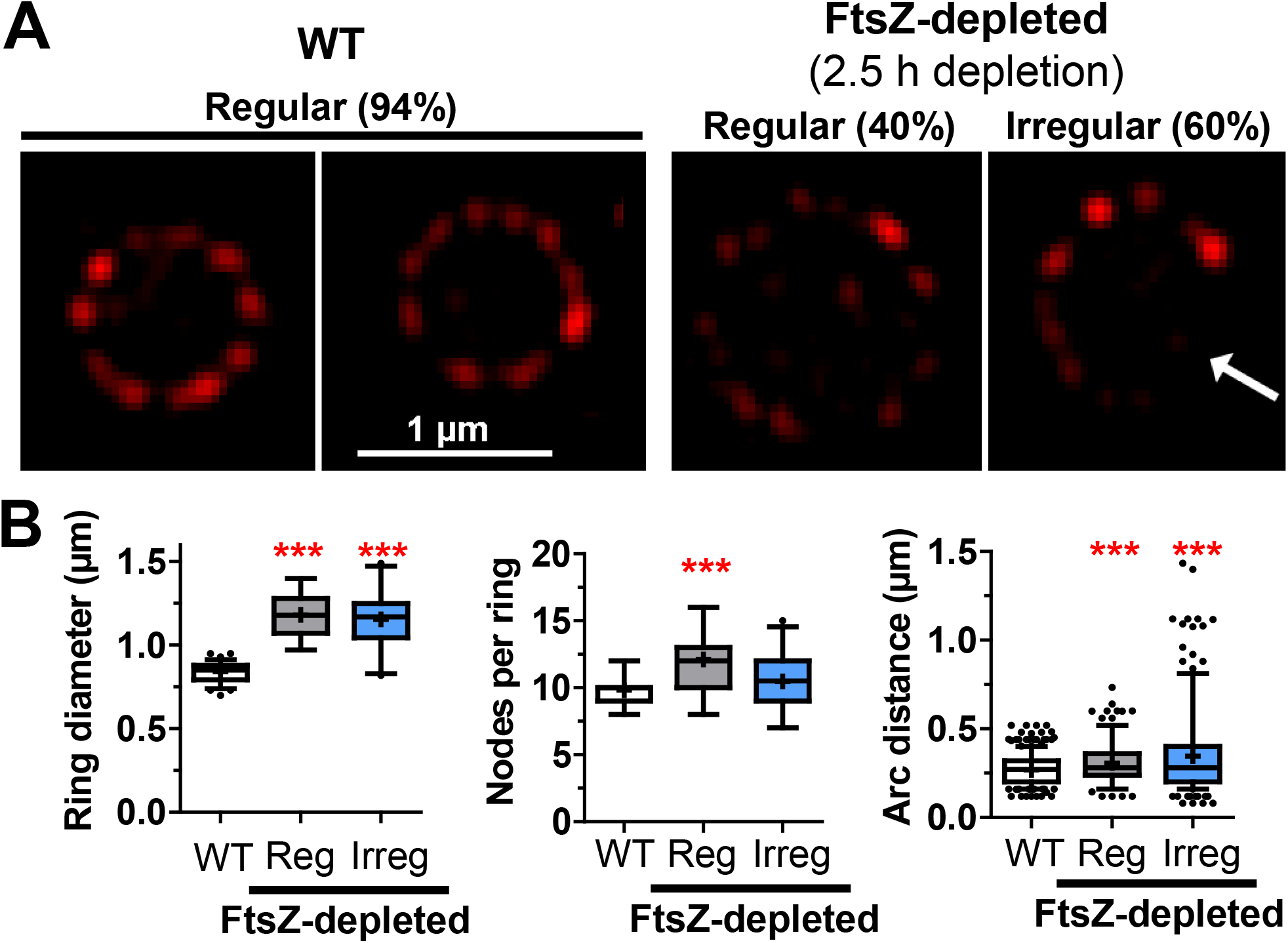
FtsZ-depleted cells have enlarged midcell diameters, and a majority of cells have irregularly spaced nodes of TP activity. IU8124 (∆*ftsZ*//PZn-*ftsZ*+) cells were grown in BHI broth containing 0.2 mM ZnCl2 to ectopically express FtsZ for 12 h prior to depletion. IU8124 cells were diluted into BHI broth without added ZnCl2 and incubated to deplete FtsZ (see *SI Appendix, Experimental Procedures*). After 2.5 h, WT (IU1945) and FtsZ-depleted cells were labeled with 45.5 μM TADA for 17 s, fixed, and prepared for vertical cell imaging by 3D-SIM (*SI Appendix, Experimental Procedures*). (*A*) Representative 3D-SIM images of nodes of TP activity in midcell rings of predivisional cells. Labeling was classified as regular (Reg) or irregular (Irreg) with gaps of varying sizes between nodes (white arrow) as described in *Results.* Percentages refer to analysis of 76 WT and 47 FtsZ-depleted cells. (*B*) Distributions of midcell ring diameters (left), nodes per ring (middle), and arc distances (right) of WT and FtsZ-depleted cells were determined as described in *SI Appendix, Experimental Procedures*. Ring diameters and nodes per ring were determined for 76 WT cells and for 19 or 28 Reg or Irreg FtsZ-depleted cells, respectively. Arc distance were determined for 746 nodes in WT cells and for 230 or 293 nodes in Reg or Irreg FtsZ-depleted cells, respectively. Graphs show median (interquartile range; whiskers, 5^th^-95^th^ percentile; +, mean). Means ± SD are compiled in Table S3. Differences in means relative to WT were determined by one-way ANOVA with Bonferonni’s multiple comparison posttest (GraphPad Prism). ***, *P*<0.001.

### The number of regularly spaced FDAA-labeled nodes is correlated with the diameter of predivisional *Spn* cells

The increase in the number of nodes in FtsZ-depleted cells with regular spacing (Fig. 5B, middle; Table S3) was suggestive of a regulatory mechanism for node placement. To explore this idea, we determined labeling patterns in five additional division and PG synthesis mutants that were pulsed with TADA for 17 s and imaged vertically (Fig. 6). Amino acid changes in FtsZ(G107S) (divisome ring organizer protein) (13), EzrA(T506I) (FtsZ-ring modulator in Gram-positive bacteria) (35), and GpsB(K96N) (regulator of the balance between sPG and pPG synthesis) (10, 36) cause temperature sensitivity (TS) at 42°C. TS mutants expressing EzrA(T506I) or GpsB(K96N) were isolated for this study (Table S1). At the semi-permissive temperature of 37°C in BHI broth, each mutant grows slower and forms cells with enlarged diameters (Fig. S4A; Table S3). The TADA-labeling pattern of each mutant was similar to that of the FtsZ-depleted strain. The majority (53% to 66%) of mutant cells show irregular nodal patterns with gaps and greater arc distances, whereas cells with regular labeling patterns contain an increased number of nodes spaced similarly to WT (Fig. 6A, 6B, and S4B; Table S3).

**Fig. 6.**
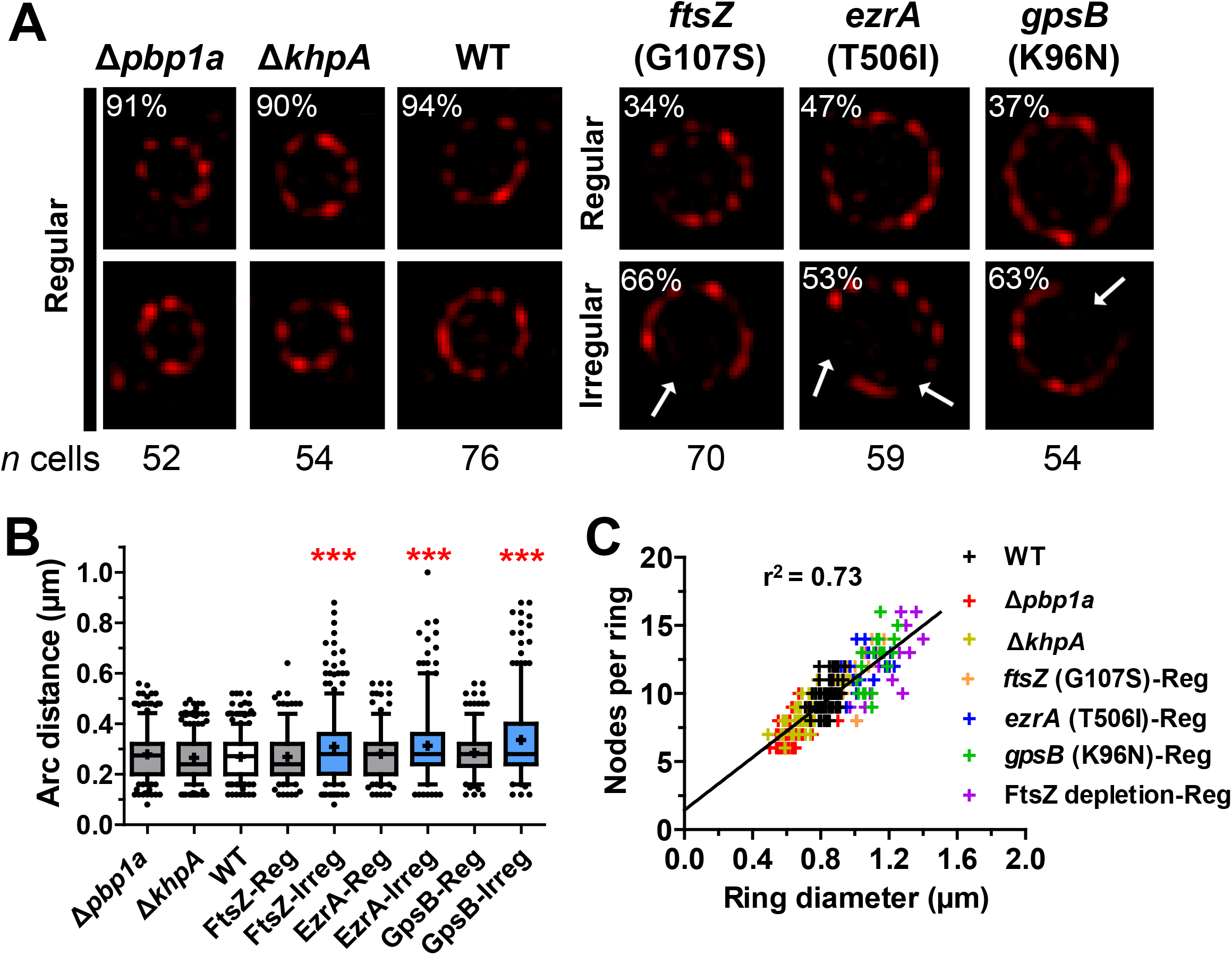
Arc distance between regularly spaced midcell TP nodes is constant in predivisional *Spn* mutants with decreased or increased diameters compared to WT. Mutant strains of *S. pneumoniae* with altered midcell diameters were labeled with 45.5 μM TADA for 17 s, fixed, and prepared for vertical cell imaging by 3D-SIM as described in *SI Appendix*, *Experimental Procedures*. Strains used were: ∆*pbp1a* (K164), ∆*khpA* (E751), WT (IU1945), *ftsZ*(G107S) (IU10612), *ezrA*(T506I) (IU11034), and *gpsB*(K96N) (IU11956) (Table S1). (*A*) Representative 3D-SIM images of TP activity in midcell rings of predivisional cells. Percentages indicate the frequency of regularly spaced nodes (Reg) or irregularly spaced nodes (Irreg) with gaps (see white arrows) for the number of cells analyzed of each strain. Irregularly spaced nodes were observed in the majority of *ftsZ*(G107S), *ezrA*(T506I) and *gpsB*(K96N) cells, whose diameters were greater than that of WT (see Fig. S4 and Table S3). (*B*) Distributions of arc distances between TP nodes of WT and mutant strains were determined as described in *SI Appendix, Experimental Procedures*. Graphs show median (interquartile range; whiskers, 5^th^-95^th^ percentile; +, mean). Means ± SD are compiled in Table S3. Cells with irregular midcell nodal patterns exhibit large SDs. Differences in means relative to WT were determined by one-way ANOVA with Bonferonni’s multiple comparison posttest (GraphPad Prism). ***, *P*<0.001. (*C*) Linear relationship between ring diameter (X-axis) versus nodes per ring (Y-axis) for WT and mutant strains with regular nodal patterns. Colored crosses represent measurements from single rings determined as described in *SI Appendix, Experimental Procedures* and shown in Fig. S4. Line of best fit and r^2^ values for the combined data set were determined using GraphPad Prism.

We extended this analysis to ∆*pbp1a* and ∆*khpA* mutants that have significantly smaller diameters than WT cells in BHI broth (Fig. S4A, Table S3). ∆*pbp1a* mutants lack aPBP1a and produce skinny, slightly elongated cells (21, 37), while ∆*khpA* mutants lack a regulatory RNA-binding protein and form smaller cells with the same aspect ratio as larger WT cells (38). The nodal pattern in the ∆*pbp1a* or ∆*khpA* mutant is regular in >90% of cells with an average arc distance similar to that of WT (Fig 6A and 6B; Table S3); consequently, the mutant cells have fewer nodes per cell (Fig. S4B). When plotted, these data show that the number of regularly spaced nodes, when present, increases linearly with ring diameter (Fig. 6C), consistent with a mechanism that maintains WT spacing between sites of PG synthesis in predivisional *Spn* cells.

### Spacing of bPBP2x and bPBP2b nodes is similar to that of FDAA nodes produced by short pulse labeling

The number and arc distance of nodes of isfGFP-bPBP2x and sfGFP-bPBP2b in the midcell rings of predivisional cells (Fig. 2A, S2A, S2B, and 7A) were measured and found to be similar to those of the FDAA nodes in WT cells (Fig. 2B and 7C; Table S3). Notably, the spacing of the TADA (red) and sfGFP-bPBP2b (green) nodes determined in the same cell is similar, but displaced in most cells (Fig. 7A), providing further support that the nodal patterns are not a microscopy artifact. To quantitate the relative distributions of these patterns, we calculated correlation coefficients for the number and spatial distribution of TADA and sfGFP-bPBP2b nodes in 48 cells (Fig. S5A and S5B). As expected by the number of nodes (Table S3), there is a positive correlation between the number of sfGFP-bPBP2b and TADA nodes (Fig. S5A). In contrast, spatial correlation coefficients were distributed around 0, indicating no strong interdependence of the positions of the TADA and sfGFP-bPBP2b nodes (Fig. S5B).

**Fig. 7.**
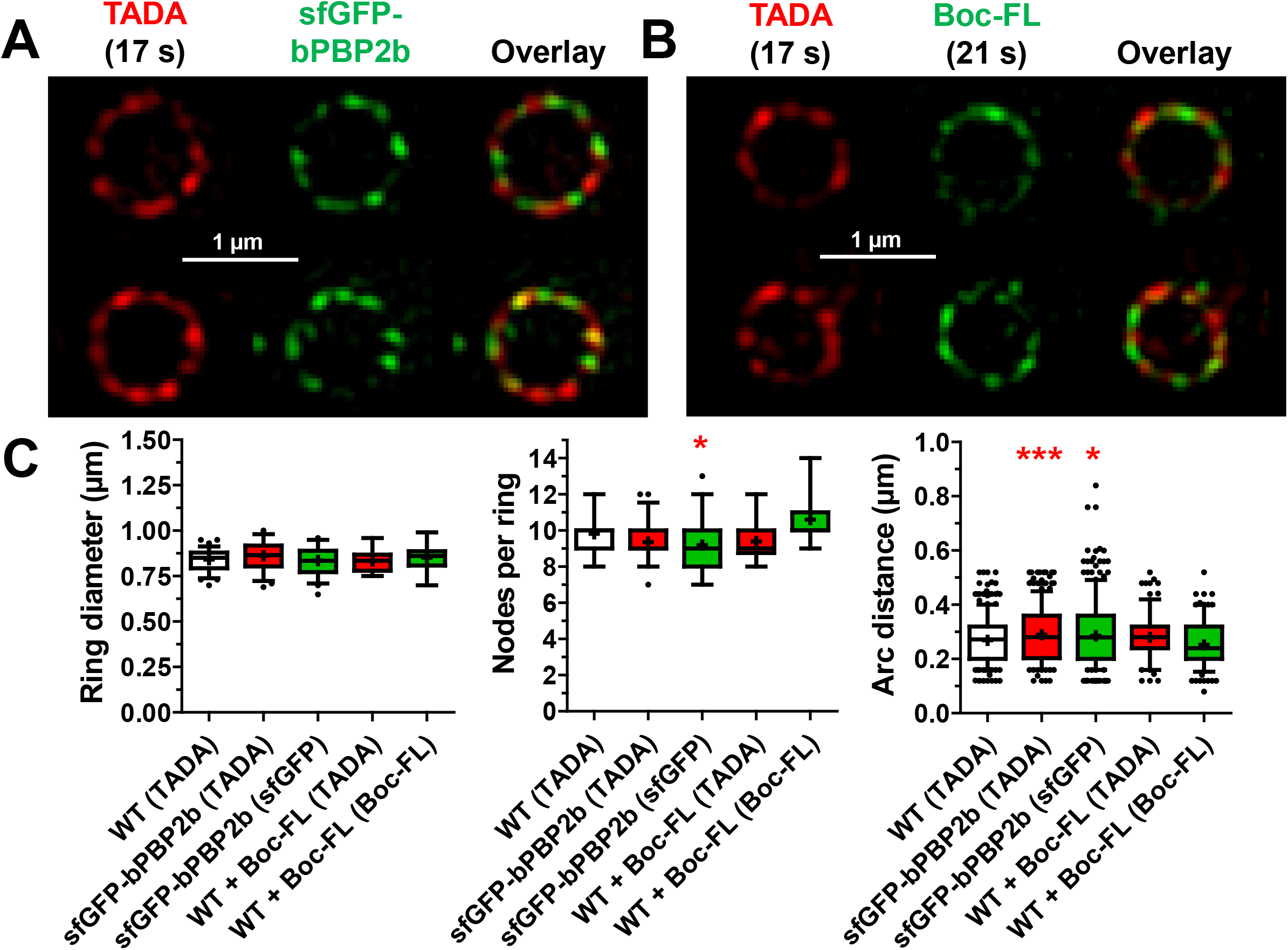
sfGFP-bPBP2b and Bocillin-FL (Boc-FL) labeling are organized in nodal patterns at the midcell of predivisional *Spn* cells. Cells expressing sfGFP-bPBP2b (IU9965) were labeled with 45.5 μM of TADA for 17 s, fixed, and prepared for vertical cell imaging by 3D-SIM as described in *SI Appendix*, *Experimental Procedures*. WT (IU1945) cells were pulse-labeled with TADA for 17 s, followed by labeling with 2 μg/mL of Boc-FL for 21 s, fixed, and prepared for imaging (see *SI, Appendix, Experimental Procedures*). (*A*) Representative images of IU9965 localizing sfGFP-bPBP2b and TADA as nodes. Each row is a separate cell. (*B*) Representative images of WT cells localizing Boc-FL and TADA as nodes. Each row is a separate cell. (*C*) Distributions of ring diameters (left), nodes per ring (middle), and arc distances (right) for WT cells labeled with TADA alone, IU9965 (sfGFP-bPBP2b) labeled with TADA, IU9965 (sfGFP-bPBP2b) not labeled with TADA, WT labeled with TADA followed by Boc-FL, and WT labeled with Boc-FL alone. Measurements were performed as described in *SI Appendix*, *Experimental Procedures*. Graphs show median (interquartile range; whiskers, 5^th^-95^th^ percentile; +, mean). Means ± SD are compiled in Table S3. Differences in means relative to WT labeled with TADA were determined by one-way ANOVA with Bonferonni’s multiple comparison posttest (GraphPad Prism). *, *P*<0.05; ***, *P*<0.001. Correlation coefficient analyses are presented in Fig. S5 and *Results*.

We performed two additional labeling protocols to gain information about these patterns. First, we performed tandem short-pulse labeling of WT cells with TADA (17 s), which labels regions of active TP activity, followed by Boc-FL (21 s), which labels all active PBPs (Fig. 7B; Table S3). Displaced two-color patterns of TADA and Boc-FL nodes were produced, similar to those of TADA-labeled sfGFP-bPBP2b cells (Fig. 7A). Again, there was a positive correlation between the number of TADA and Boc-FL nodes (Fig. S5C), but no spatial correlation of the positions of the two kinds of nodes in 18 cells (Fig. S5D). The regular nodal pattern after Boc-FL pulse-labeling pattern did not match what would be expected from an earlier study done at long labeling times (39). We reprised this earlier study and obtained different results (*Appendix SI, Additional data*; Fig. S6), consistent with those reported here (Fig. 7B). Last, we tandemly pulse labeled WT cells with three different colors of FDAAs (green, then red, then blue) (Fig. S7). We again observed regular, displaced nodal patterns for each color of FDAA probe (arrows, Fig. S7), supporting the conclusion that there is a distributive, regular nodal pattern of PBP localization and TP activity at the midcell ring of predivisional *Spn* cells.

### FDAA labeling is organized in nodes in other ovoid-shaped bacterial species

To determine if this regular, organized pattern of TP activity by PBPs is present in other ovococcal species, we labeled *Streptococcus mitis* and *Enterococcus faecalis* with short (17 s) pulses of TADA and imaged vertically oriented cells by 3D-SIM (Fig. 8). *S. mitis* is a close evolutionary relative that exchanges DNA with *Spn* by natural competence, while *E. faecalis* is more evolutionarily distant in the same *Lactobacillale* order (40–42). Both species demonstrated a pattern of TADA-labeled nodes at midcell with an average arc distance indistinguishable from that of WT *Spn* (Fig. 8B; Table S3). Thus, the regular organized pattern of PBP TP activity in predivisional cells is widely distributed in ovococcal species.

**Fig. 8.**
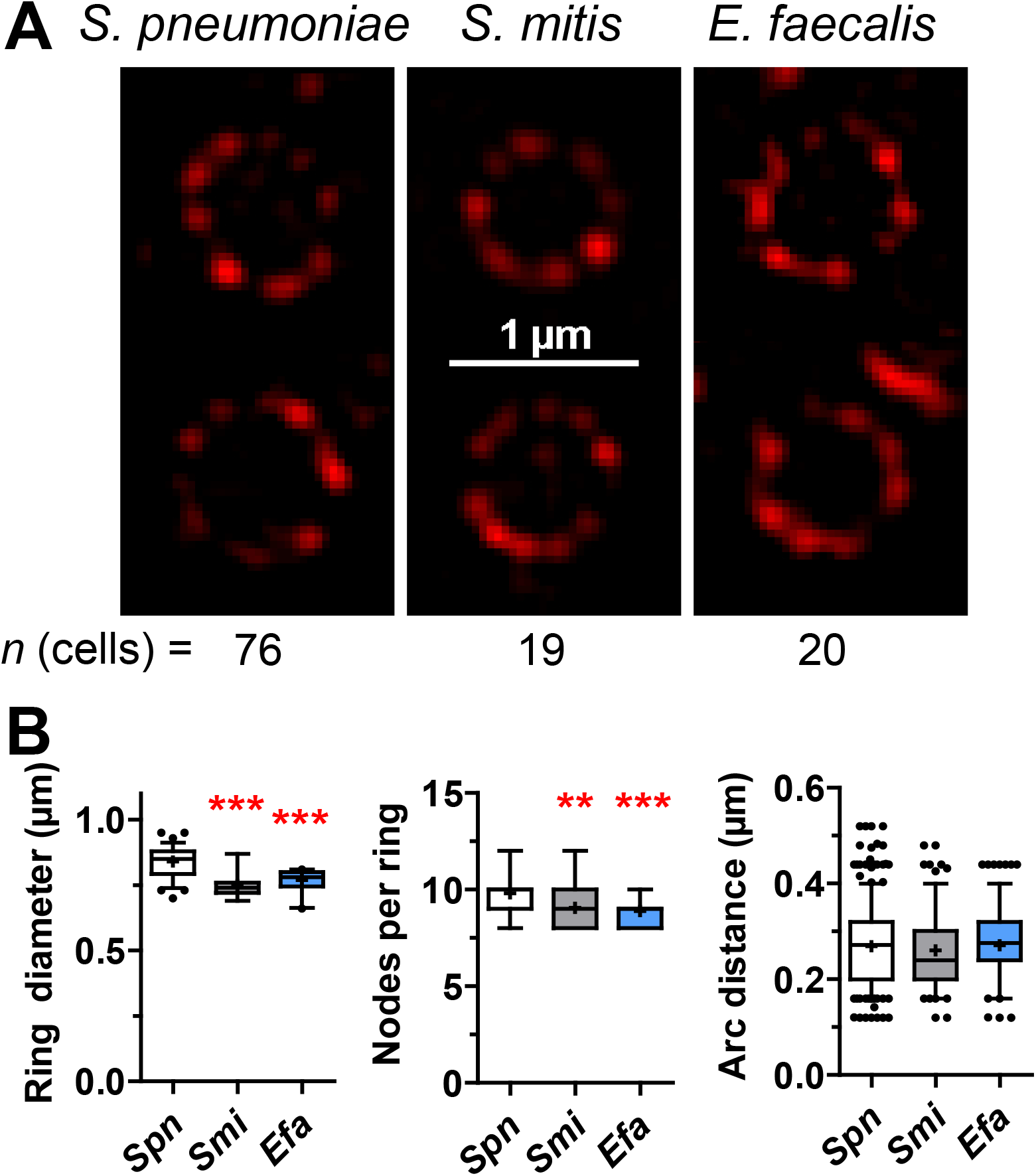
The nodal pattern of regularly spaced TP activity at the midcell of predivisional cells is conserved in other ovoid-shaped bacterial species. *S. pneumoniae* (*Spn*, IU1945), *S. mitis* (*Smi*, ATCC 49456) and *E. faecalis* (*Efa*, ATCC 51299) cells were labeled with 45.5 μM TADA for 17 s, fixed, and prepared for vertical cell imaging by 3D-SIM (see *SI Appendix*, *Experimental Procedures*). (A) Representative images of two predivisional cells pulse-labeled with TADA are shown for *Spn* (left), *S. mitis* (middle), and *E. faecalis* (right) from the total number of cells analyzed (bottom). (B) Distributions of ring diameters (left), nodes per ring (middle), and arc distances of ovoid-shaped cells. Measurements were performed as described in *SI Appendix*, *Experimental Procedures*. Graphs show median (interquartile range; whiskers, 5^th^-95^th^ percentile; +, mean). Means ± SD are compiled in Table S3. Differences in means relative to *Spn* were determined by one-way ANOVA with Bonferonni’s multiple comparison posttest (GraphPad Prism).**, *P*<0.01 and ***, *P*<0.001.

## DISCUSSION

Whether the sPG and pPG synthesis machines of *Spn* are distributed contiguously in a single midcell ring (“plum pudding” model) or separate from each other during septal constriction has been a point of contention (11, 43–45). The demonstration of concentric intermediate rings of TP activity reported here provides strong support for the separation hypothesis (Fig. 1D and 9A). Localization of bPBP2x to the inner ring, which corresponds to the constricting leading edge of the septal annulus, further supports closure driven by PG synthesis by the bPBP2x:FtsW complex (13, 18). Some bPBP2x remains in the outer ring of mid-to-late divisional cells when iHT-bPBP2x is expressed at near WT level in C+Y medium (Fig. S2E) or when WT bPBP2x is itself labeled with the 7FL probe (Fig. S2F) (25). Thus, active bPBP2x is both at the constricting leading edge of the septal annulus and at its outer edge, suggesting expansion of the annulus in both places. The constricting FtsZ ring also tracks with sPG synthesis at the leading edge of the septal annulus ring, implying that FtsZ is not detectable in the outer pPG synthesis ring (Fig. S2H).

In contrast, bPBP2b expressed at 12% or at WT levels primarily localizes to the outer pPG synthesis ring (Fig. 2A, S2B, S2D, and S3C). These results indicate that only about 12% of bPBP2b cellular amount is sufficient for normal growth and *Spn* cell morphology (Fig. S3A and S3B), implying that bPBP2b is in excess in cells in this culture condition. An implication of confinement of bPBP2b to the outer midcell ring is that the presumed circumferential movement of the bPBP2b:RodA complex during pPG synthesis is guided by a structure that lacks FtsZ filaments/bundles. *Spn* does not encode the actin-like MreB protein that mediates PG elongation of rod-shaped bacteria (4), and determination of *Spn* elongasome composition and organization is an area of active research.

FtsX is also confined to the outer pPG synthesis ring (Fig. S2G). FtsX is an essential, polytopic membrane protein that forms a complex with the cytoplasmic FtsE ATPase and the extracellular PcsB PG hydrolase in *Spn* (28–32). Depletion of essential FtsX, FtsE, or PcsB results in chains of spherical cells, similar to those caused by depletion of bPBP2b (11, 28, 30, 31, 33, 46). An FtsX’-isfGFP-FtsX’ fusion was detected in the outer PG synthesis ring, but not in the inner septal ring (Fig. S2G), consistent with a role for FtsEX:PcsB in pPG synthesis, analogous to that played by FtsEX:CwlO in sidewall PG synthesis in *Bacillus subtilis* (*Bsu*) (47, 48). Thus, *Spn* FtsX is in proximity to FtsZ and FtsA in the nascent divisome in early predivisional cells (Fig. S2G); however, in contrast to *E. coli* FtsX (49), as division proceeds, *Spn* FtsX physically separates from FtsZ and FtsA, which are at the leading edge of the septal annulus (Fig. S2H). This result raises the possibility that a PG remodeling hydrolase other than FtsEX:PcsB mediates sPG synthesis. Based on these examples, dual labeling of vertical cells can be applied to assign other PG synthesis, divisome, and regulatory proteins to the sPG or pPG synthesis machines of *Spn*. This approach may also explicate PG stress responses, such as possible roles of Class A PBPs in imparting resistance to an exogenously added PG hydrolase following exposure of *Spn* laboratory strain R6 to a β-lactam antibiotic that inhibits bPBP2x (and DacA (PBP3)) (14).

We noted that bPBP2x and bPBP2b are distributed in regular nodal patterns at the equators of predivisional cells (Fig. 2, 7A, and S2 A-F). As the inner ring constricts, this nodal pattern becomes more compact and difficult to resolve (Fig. 2A); hence, we confined this study to the large equators of predivisional cells (Fig. 3A). By shortening the FDAA pulse time down from 2.5 min to ≈17 s, we observed that TP activity also is distributed in a regular nodal pattern with ≈10 nodes separated by an average arc length of 0.27 ± 0.07 μm in WT cells (Fig. 3B). A comparable nodal pattern of FDAA pulse labeling was detected in *S. mitis* and *E. faecalis* (Fig. 8), indicating a common organization of TP activity in predivisional ovococcal cells.

Several controls support the conclusion that the nodal pattern was not caused by processing steps required to generate structured illumination images (see *Results)*. Most importantly, the regular nodal pattern of FDAA labeling was disrupted by gaps in a majority of cells depleted for FtsZ (Fig. 5) or in temperature-sensitive *ftsZ*(G107S), *ezrA*(T506I), and *gpsB*(K96N) mutants grown at the semi-permissive temperature of 37°C (Fig. 6). Each of these growing TS mutants formed enlarged cells with increased diameters compared to WT (Fig. S4). Between 34%-47% of these larger mutant cells still showed regular nodal patterns of FDAA labeling with more nodes per ring spaced at the WT arc distance (Fig. 6). Conversely, ∆*pbp1a* or ∆*khpA* mutants have smaller diameters than WT with fewer FDAA nodes spaced at the WT arc distance (Fig. 6). Together, these data are consistent with a mechanism that maintains a regular spacing of PG synthesis in predivisional cells, irrespective of equator diameter (Fig. 6C and 9B).

The average arc distance was similar between nodes of FDAA labeling, isfGFP-bPBP2x, isfGFP-bPBP2b, and Boc-FL (Fig. 2, 3, and 7). However, the positioning of nodes in two-color FDAA/sfGFP-bPBP2b or FDAA/Boc-FL experiments showed low correlation (Fig. S5), as did nodes sequentially pulse-labeled with three different colors of FDAAs (Fig. S7). The displacement of FDAA nodes relative to bPBP2b or Boc-FL, may reflect the time (≈1 min) that it takes to wash away unincorporated FDAA. It is also possible that PBPs other than bPBP2b are active in predivisional cells. Altogether, these results indicate that the placement of PBPs and TP activity is not fixed at single positions on equators of predivisional cells, but rather, is dynamic and distributive.

The localization of PG synthesis and PBPs in early divisional cells varies among bacterial species that use different modes of septum formation and division. Similar to the patterns reported here for *Spn* and other ovococcal species (Fig. 2, 8, and S2), labeling *Eco* for short pulses, but not for long times, with an FDAA results in nodal (also called punctate) patterns, which were not quantitated at high resolution (50). Consistent with nodal pattern formation, high-resolution microscopy of vertically oriented *Eco* cells revealed a pattern of “discrete densities” of mCit-FtsI (bPBP3) distributed around the entire septal ring of dividing *Eco* cells (51).

Different FDAA pulse-labeling patterns were reported for *S. aureus* (*Sau*) and *Bsu* compared to those in *Spn* and *Eco*. Discrete foci of PG synthesis were not observed in *Sau* cells labeled with short FDAA pulses and viewed by 3D-SIM, in an experiment similar to Figure 3, or labeled with a pulse of other D-amino acid probes and viewed by localization microscopy (52). However, while not commented upon, SIM images of sfGFP-PBP1 in some vertically oriented *Sau* cells do appear nodal around closing septal rings (53), and heterogenous localization of *Sau* GFP-PBP2 has also been reported (54). Labeling *Bsu* with short, sequential pulses of two colors of FDAAs or an FDAA followed by Boc-FL (55) gave a different pattern from the regular nodal pattern in *Spn* (Fig. 7 and S7). At the lower resolution of rotated 3D-SIM images, sequential labeling with two colors of FDAAs gives a pattern suggestive of a limited number of PG synthesis complexes moving in both directions around the septum. Labeling *Bsu* with an FDAA followed by Boc-FL again suggests a surprisingly limited number of active PBP complexes labeled by Boc-FL adjacent to newly synthesized PG marked by the FDAA (55). We conclude that nodal PG synthesis at septa occurs in different patterns in some bacteria and is apparently absent in others, possibly reflecting different mechanisms of sPG synthesis and cell separation. In this regard, *Spn* and *Eco* simultaneously close division septa while separating daughter cells (13, 50), whereas septa formation by PG synthesis occurs before cell separation in *Sau* and *Bsu* (52, 53, 55).

The mechanism that causes the regular nodal distribution of PBPs and their TP activity in predivisional cells of *Spn* remains to be determined (Fig. 9B). Recent notable studies have demonstrated heterogeneous supramolecular localization of membrane proteins generally and at septa of *Sau* cells, including interacting enzymes that catalyze phospholipid biosynthesis and the MreD regulator of PG synthesis (56, 57). This heterogenous localization results in punctate patterns of these proteins at division septa, resembling the localization of *Spn* PBPs reported here (Fig. 2 and S2). Polytopic MreD plays a role in supramolecular localization of the phospholipid biosynthesis enzymes, and heterogeneous punctate patterns are lost upon protein overexpression (57). A favored explanation for punctate supramolecular organization is that membrane protein complexes distort local curvature on membranes and thereby perturb diffusion, resulting in distribution patterns for the majority of membrane proteins (57). Whether membrane protein distribution, the size and composition of PG synthesis complexes, an unknown scaffolding complex, or a combination of these mechanisms causes the nodal distribution of PBPs and TP activity in predivisional *Spn* cells remains to be determined.

**Fig. 9.**
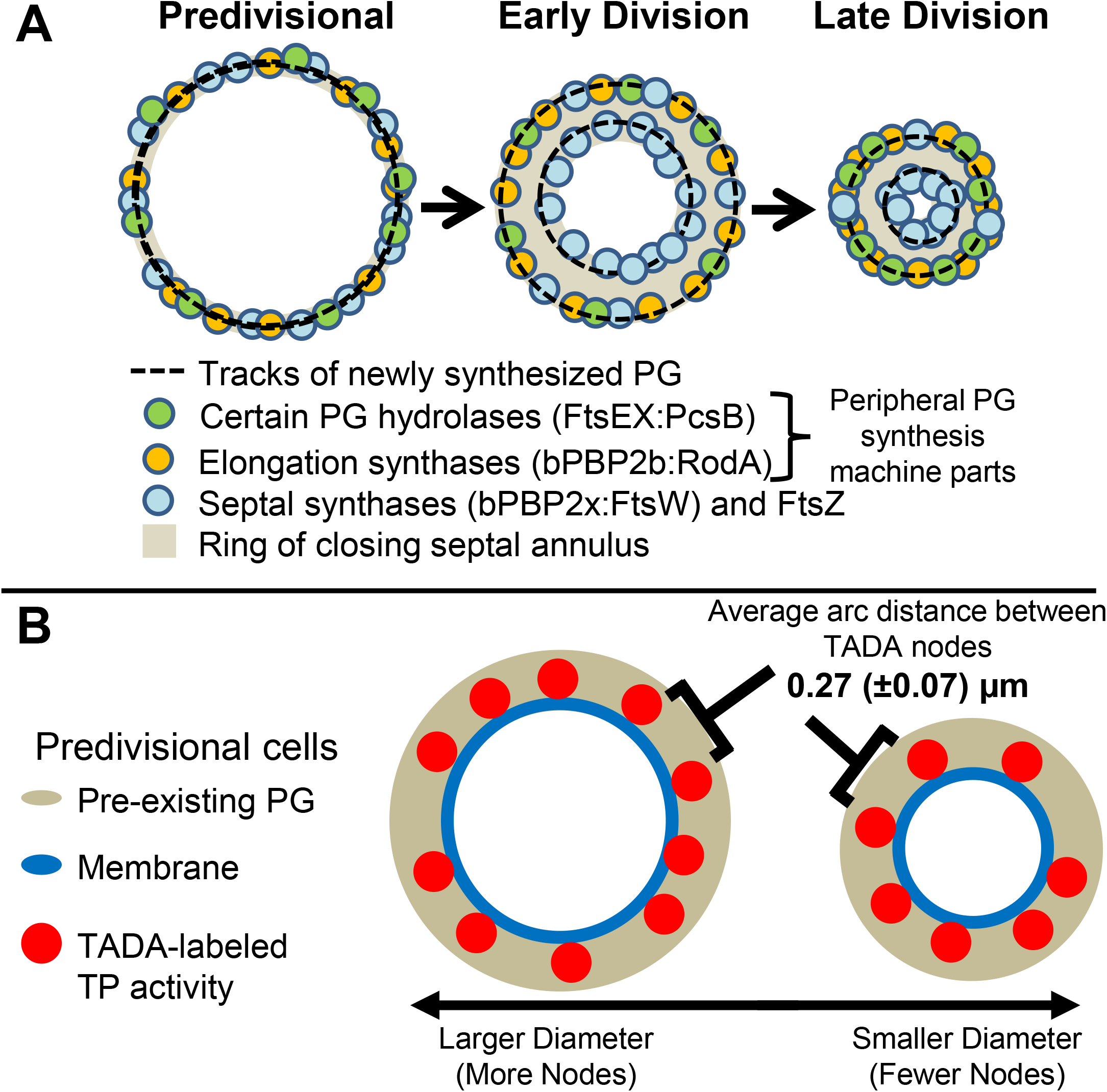
Summary diagram of the localization PG synthesis in *Spn* based on results in this paper. (*A*) Separation of sPG and pPG (elongasome) machines at the midcell of dividing *Spn* cells. Cell division begins at the midcell equator of newly divided predivisional cells in FtsZ-organized divisome rings containing the components of the sPG and pPG synthesis machines. Early in cell division the septal annulus forms and begins to close. The leading edge of the closing septal annulus separates the sPG machine from the pPG synthesis machine that remains at the outer edge of the annulus and elongates the PG outward from the midcell. The sPG machine contains bPBP2x TP, its partner FtsW GT, and other components, while the pPG synthesis machine is made up of bPBP2b TP and its partner RodA GT, other “Rod” complex proteins, and the FtsEX:PcsB remodeling PG hydrolase. The constricting FtsZ ring tracks with the leading edge of the septal annulus, such that the outer peripheral ring is not organized by FtsZ beyond the predivisional stage. Later in division, FtsZ remaining at the septum migrates to the developing equatorial rings in daughter cells. The inner sPG synthesis machine constricts into a dot surrounded by the closing outer pPG synthesis ring, which eventually constricts and merges with the sPG dot as the new cell pole is completed and the PG synthesis enzymes migrate to the new equatorial rings in daughter cells (see Fig. 1D also). (*B*) In predivisional cells, TP activity, sPG synthesis complexes, and pPG complexes are organized into a pattern of regularly spaced nodes (red circles). The placement of PBPs and TP activity is not fixed at single positions on equators of early divisional cells, but rather, is dynamic and distributive, likely driven by PG synthesis itself (13). Aggregate movements of these heterogenous nodal complexes with time would account for the distributive, non-correlated nodal patterns observed in two-color labeling experiments. A constant distance (≈0.27 μm) is maintained between nodes of PBP TP activity and between the PBPs themselves. Midcell rings of predivisional cells with smaller or larger diameters contain fewer or more nodes, respectively, than WT (≈ 10 nodes).

Finally, single molecules of bPBP2x and its interacting partner FtsW move circumferentially around the septa of dividing pneumococcal cells in either direction at ≈21 ± 8 (SD) nm/s (13). Based on data in that paper, runs extend for ≈28 ± 15 (SD) s (n = 106 cells). In contrast, bPBP2x and FtsW not bound to septa move diffusively throughout the cell membrane (13, 58). In *Spn*, movement of the bPBP2x:FtsW complex at septa is driven by PG synthesis itself and not by treadmilling movement of FtsZ filaments/bundles (13). In the static images here, we do not precisely know how many molecules form the bPBP and FDAA-labeled nodes; but, a reasonable assumption is that each node consists of multiple PBPs and regions of PG labeling. With this in mind, circumferentially moving bPBP2x:FtsW complexes likely move linearly in both directions across nodal regions driven by the PG synthesis echoed by FDAA labeling. Aggregate movements of these heterogenous nodal complexes with time would then account for the distributive, non-correlated labeling patterns in two-color pulse labeling experiments (Fig. 7B and S7). This and other hypotheses about the composition, organization, and dynamic movement of these nodal PG synthesis complexes in *Spn* and other ovococcal bacteria await future testing.

## EXPERIMENTAL PROCEDURES

Detailed experimental procedures are described in *SI Appendix, Experimental Procedures*, including bacterial strains (Tables S1 and S2) and growth conditions; growth curve analysis; 2D-epifluorescence and phase-contrast microscopy of *Spn* cells; saturating labeling with Boc-FL and Ceph-CT; FDAAs used; quantitative western blotting of relative cellular amounts of WT and fusion proteins; localization in vertical *Spn* cells of the following probes after the indicated labeling times: HADA (≈100 min) then TADA (2.5 min), TADA (2.5 min) in cells expressing isfGFP-bPBP2x, sfGFP-bPBP2b, or FtsX’-isfGFP-FtsX’, HADA (2.5 min) in cells expressing iHT-bPBP2x or iHT-bPBP2b, TADA (17 s); localization of bPBP2x with 7FL in WT *Spn* and a ∆*pbp1b* mutant; localization in vertical *Spn* cells of the following probes after the indicated labeling times: TADA (17 s), TADA (17 s) followed by Boc-FL (7 s), and BADA (40 s) followed by TADA (40 s) followed by HADA (40 s); image acquisition by 3D-SIM; analysis of FDAA node distributions using a custom MATLAB GUI; widefield fluorescence microscopy; and bPBP2x and bPBP2b protein purification and antibody production.

## ACKNOWLEDGEMENTS

We thank laboratory members and Erkin Kuru (Harvard Med Sch), Yves Brun (Université de Montréal), Seamus Holden (Newcastle Univ), and Jie Xiao (Johns Hopkins) for discussions and suggestions; Jim Powers (IUB) for help with microscopy, Luke Lavis (Janelia Lab) for Fluor JF549; Dalia Denapaite (Trento Univ), Reinhold Brückner, and Regine Hakenbeck (Kaiserlautern Univ) for anti-bPBP2x antibody; and Suzanne Walker and David Z. Rudner (Harvard Med Sch) for supporting preparation of anti-bPBP2x and anti-bPBP2b antibodies. This work was supported by NIH Grants RO1GM113172 (to M.S.V. and M.E.W.); RO1GM128439 (to E.E.C. and M.E.W.); R35GM131767 (to M.E.W.); R35GM136365 (to M.S.V), RO1AI148752 (to S. Walker), RO1AI139083 (To D. Z. Rudner), National Science Foundation Grant MCB1615907 (to S.L.S.); NIH Predoctoral Quantitative and Chemical Biology Training Grant T32 GM109825 (to A.J.P.); NIH Predoctoral Grant F31AI138430 (to M.M.L.); and NIH Equipment Grant S10OD024988 to the Indiana University Bloomington (IUB) Light Microscopy Imaging Center.

