## Supplemental Tables, Figures, and Data for "Organization of Peptidoglycan Synthesis in Nodes and Separate Rings at Different Stages of Cell Division of *Streptococcus pneumoniae*"

Running title: Microscopy of Vertically Oriented Pneumococcal Cells

[†]Contributed equally to this work. Author order was determined by random draw.

**SI APPENDIX, EXPERIMENTAL PROCEDURES**

**SI APPENDIX, ADDITIONAL DATA**

**SI APPENDIX, TABLES:** Tables S1-S3

**SI APPENDIX, SUPPLEMENTAL FIGURES AND LEGENDS**: Fig. S1-S7

**SI APPENDIX, REFERENCES**

[#]Corresponding author:
Malcolm E. Winkler
Department of Biology
Indiana University Bloomington
1001 East Third Street
Bloomington, Indiana USA 47405


### SI APPENDIX, EXPERIMENTAL PROCEDURES

**Bacterial strains and growth.** *S. pneumoniae* (*Spn*) strains used in this study were constructed in strains IU1824 and IU1945, which are unencapsulated (Δ*cps*) derivatives of serotype 2 strain D39W (*SI, Appendix*, Table S1). Strains were constructed by transforming linear DNA amplicons into competent cells and culturing transformants on trypticase soy agar II plates (BD BBL, 221261) containing 5% (v/v) defibrinated sheep blood as described previously (1), except when noted (see *SI Appendix,* Table S1 footnotes). Primers used for the synthesis of amplicons are listed in *SI Appendix,* Table S2.

Strains were inoculated from frozen glycerol stocks into 5 mL of brain heart infusion (BHI; BD Bacto, 237500) broth containing added $ZnCl_2$ and $MnSO_4$ where indicated. Serial dilutions were prepared and incubated for 12-16 h as overnight cultures at 37°C in an atmosphere of 5% $CO_2$. Extent of growth of liquid cultures was determined by $OD_{620}$ as described previously (2). Overnight cultures still in exponential phase ($OD_{620}$ ≈0.05-0.4) were diluted in parallel to $OD_{620}$ ≈0.003-0.005 in 5 mL of fresh BHI broth, containing added $ZnCl_2$ and $MnSO_4$ where indicated. Cultures were incubated at 37°C in 5% $CO_2$, and $OD_{620}$ readings were taken every 45-60 min until cultures reached stationary phase.

**2D-epifluorescence microscopy (2D-EFM) and phase-contrast microscopy.** *Spn* cells were prepared for microscopy as described below in sections describing labeling. Cells were viewed on a Nikon Eclipse E-400 epifluorescence phase-contrast microscope using a Nikon Intensilight C-HGFI epifluorescence illuminator. Images were captured using a CoolSNAP HQ2 charge-coupled device (CCD) camera (Photometrics) and processed with Nikon NIS-Elements BR imaging software. sfGFP and isfGFP images were collected using the FITC-HYQ filter (Ex 460-500 nm; Em 560 nm) with a 1 s exposure time. Halo-Tag (HT) images were collected using the Texas Red-HYQ filter (Ex 532-587 nm; Em 650 nm) with a 1 s exposure time. Additional image processing was completed using FIJI (3).

**Fluorescent D-amino acids (FDAAs) used.** FDAAs were synthesized as reported in (4), with the following change: TADA was synthesized as reported for TDL, except that Boc-D-DAP-OH (*N*-alpha-t-butyloxycarbonyl-D-2,3-diaminopropionic acid) was used in place of Boc-D-Lys-OH (*N*-alpha-t-butyloxycarbonyl-D-lysine). Working solutions of HADA and TADA in BHI broth were diluted from 500 mM stocks in DMSO, which were stored at -20°C and shielded from light.

**Localization of HADA (≈100 min labeling), then TADA (2.5 min labeling) in vertically oriented *Spn* cells.** Cells from overnight cultures were diluted to $OD_{620}$ ≈0.02 in 2 mL of fresh BHI broth containing HADA (125 µM final). At $OD_{620}$ ≈0.22, 575 µL of cultures was centrifuged at 16,100 × *g* for 5 min at room temperature, and pellets were resuspended in 250 µL of BHI broth containing TADA (125 µM final). Cells were incubated at 37°C for 2.5 min, chilled on dry ice for 20 s, and centrifuged for 2.5 min at 16,100 × *g* at 4°C. Cell pellets were centrifuged and washed twice with 500 µL ice-cold PBS, then centrifuged a third time and resuspended in 500 µL of 4% (v/v) paraformaldehyde and incubated in the dark for 15 min at room temperature, followed by 45 min on ice in the dark.

Fixed cells were centrifuged, and pellets were resuspended in 100 µL of ice-cold GTE buffer (50 mM glucose, 20 mM Tris-HCl, pH 7.5, 1 mM EDTA). For imaging, cells were centrifuged for 5 min at 16,100 × *g* at 4°C to remove the GTE buffer and centrifuged once more to remove residual GTE buffer with a P20 pipette. Excess liquid was allowed to evaporate for approximately 1 min before pellets were resuspended in 3.0 µL of Vectashield Hardset Antifade (Vector Laboratories, H-1400) with vortexing. 1.2 µL of resuspended cells was pipetted onto a 12 mm/1.5 round coverslip (EMS, 72230-01), and a microscope slide was carefully placed on top. The slide was incubated coverslip side down at room temperature in the dark for 15 min, and then viewed by 3-Dimensional Structured Illumination Microscopy (3D-SIM). Exposure time and %T settings to acquire images were 5 ms and 50% for HADA and for TADA.

**Quantitative western blotting of relative cellular amounts of fusion proteins.** The amounts of sfGFP- and HT-tagged fusion proteins present relative to WT untagged protein were determined by quantitative western blotting with antibodies to native proteins. Cells from overnight cultures were diluted to $OD_{620}$ ≈0.005 in 5 mL of fresh BHI broth. At $OD_{620}$ ≈ 0.15-0.2, 2 mL of culture was collected and cells were centrifuged for 5 min at 16,100 × *g* at 4°C, washed in 2 mL of ice-cold PBS, centrifuged, and the supernatant removed. Cell pellets were placed on dry ice for 15 min, and thawed at room temperature for 5 min. Pellets were resuspended in 80 µL SEDS lysis buffer (0.1% (v/v) deoxycholate, 150 mM NaCl, 0.2% (v/v) SDS, 15 mM EDTA, pH 8.0), vortexed vigorously, and incubated at 37°C with shaking at 300 rpm for 15 min, with brief vortex mixing every 5 min. Protein concentrations of cell lysates were determined using the DC protein assay kit (Bio-Rad, 5000116) with a standard curve of 0.1-1.3 mg/mL of BSA as described by the manufacturer's instructions. Samples were diluted 1:1 with 2x Laemmli sample buffer (Bio-Rad, 161-0737) containing 5% (v/v) β-mercaptoethanol and incubated at 95°C for 10 min. Cell lysates were loaded onto a 10% SDS-PAGE gel (Bio-Rad, 456-1035), run for 1 h at 150 V and transferred onto a nitrocellulose membrane for 1.5 h at 350 mA. Blots were blocked in 6 mL PBS-Tw (PBS with 0.1% (w/v) Tween 20 (Sigma, P1379-500ML)) containing 50 mg/mL skim milk powder (BD Difco, 232100) for 20 min, then rinsed briefly twice in 5 mL PBS-Tw. Blots were incubated for 1 h in 10 mL PBS-Tw with either rabbit anti-bPBP2b or rabbit anti-bPBP2x antibodies (1:10,000 dilution). Blots were rinsed briefly twice in 5 mL PBS-Tw, then washed in 5 mL PBS-Tw for 15 min. Blots were incubated for 1 h in 10 mL PBS-Tw with ECL anti-rabbit IgG horseradish peroxidase-linked whole antibody (1:10,000 dilution; GE Healthcare, NA93AV), then successively washed for 5, 5, 15, 5, and 5 min in fresh PBS-Tw. Blots were incubated for 1 min with Amersham ECL western blotting detection reagent (GE Healthcare, RPN2106), and imaged with an IVIS imaging system (Xenogen) using a 1 min exposure time, as described before (5).

For experiments to determine FtsX′-isfGFP-FtsX′ levels, the following changes were made, as described in (6). Briefly, cells from overnight cultures were diluted to $OD_{620}$ ≈0.002 in 25 mL of fresh BHI broth. At $OD_{620}$ ≈ 0.15-0.2, cultures were centrifuged for 10 min at 8,000 × *g* at 4°C, and pellets were resuspended in 1.5 mL ice-cold buffer containing 20 mM potassium phosphate, pH 7.5 and 140 mM NaCl. Samples were transferred to 2 mL Lysing Matrix B Fast Prep tubes (MP Biomedicals 116911050) and lysed at 4°C in a FastPrep-24 (MP Biomedicals) using 3 runs of 40 s at 6.0 m/s in a 24x2 adaptor. Samples were placed on ice, then centrifuged for 1 min at 14,000 × *g* at 4°C. 1.2 mL supernatant and 1 mL buffer were placed in a 3.2 mL polypropylene ultracentrifuge tube (Beckman Coulter, 362333), and tubes were balanced with buffer and centrifuged for 45 min at 100,000 × *g* at 4°C in a Beckman Coulter Optima Max-XP ultracentrifuge, using a TLA-100 fixed-angle rotor. Pellets were resuspended in 250 µL of 20 mM potassium phosphate, pH 7.5, 140 mM NaCl, and 0.04% (w/v) n-dodecyl β-D-maltoside (Pierce, 89903). Protein assays, gel electrophoresis, transfer to nitrocellulose membranes, and blocking procedures were performed as described above. Blots were incubated for 1 h in 10 mL PBS-Tw containing 100 mg BSA with polyclonal rabbit anti-FtsX (1:500 dilution) (6). Secondary antibody labeling, chemiluminescent detection, and imaging were performed as described above.

To construct a standard curve, western blots of three or four different amounts of the same extract sample (ranging from 1.25 to 12.0 µg of total protein) were performed as described above, and the chemiluminescent signal of protein bands was plotted against the amount of protein loaded. Linear regression analysis (GraphPad Prism) was used to determine the linear signal range for each antibody used. For samples determined to be in the linear range, relative protein amounts were determined by dividing each signal by the signal of the untagged WT protein (≡1).

**Localization of TADA (2.5 min label) and sfGFP-tagged proteins in vertically oriented *Spn* cells.** For strains IU1824 (WT), IU11157 (*isfgfp-pbp2x*), IU9965 (*sfgfp-pbp2b*), and IU16121 (*ftsX'-isfgfp-ftsX'*), cells from overnight cultures were diluted to $OD_{620}$ ≈0.003 in 5 mL of fresh BHI broth. At $OD_{620}$ ≈0.23, 600 µL of culture was transferred to a 0.22 µm microcentrifuge tube filter (Corning Costar Spin-X, CLS8160) and the media was removed by applying vacuum to the filter with a vacuum pump. The filter was incubated with 250 µL of BHI broth at 37°C for 3 s, which was filtered away quickly with vacuum. 500 µL PBS at 37°C was added to the filter and immediately filtered away. The filter was removed from the vacuum pump and 250 µL of BHI broth containing TADA (125 µM final) was pipetted up and down approximately 10 times to resuspend the cells, which were incubated at 37°C in the dark for 2.5 m. After incubation, the filter was reattached to the vacuum pump and the media was filtered away. 500 µL of PBS at room temperature was added to the filter and immediately filtered away. This wash step was repeated, then the filter was removed from the vacuum pump. 600 µL of 4% (v/v) paraformaldehyde was added to the filter and mixed by pipetting up and down approximately 10 times to resuspend the cells, which were transferred to a 2 mL tube (Eppendorf 022431102) and incubated in the dark for 15 min at room temperature, then 45 min on ice in the dark. After fixing, cells were centrifuged for 5 min at 16,100 × *g* at 4°C and washed twice with 600 µL ice-cold PBS. Cells were then washed in 200 µL of ice-cold GTE buffer. Cells were prepared and imaged as described above. Exposure times and %T settings were 5 ms and 50% for TADA and for sfGFP.

**Localization of HADA (2.5 min label) and HT-tagged proteins in vertically oriented *Spn* cells.** For strains IU1824 (WT), IU14927 (*iht-pbp2x*), IU15928 (*iht-pbp2b*) and IU16553 (*iht-pbp2b*//$P_{Zn}$-*iht-pbp2b*), cells from overnight cultures were diluted to $OD_{620}$ ≈0.003 in 5 mL of fresh BHI broth (for IU16553, BHI broth was supplemented with 0.3 mM $ZnCl_2$ and 0.03 mM $MnSO_4$ throughout). At $OD_{620}$ ≈0.07, JF549 Halo-Tag ligand (400 µM stock in DMSO) was added to 600 µL culture to a final concentration of 500 nM. Cultures were incubated at 37°C in 5% $CO_2$ in the dark for 30 m, with brief vortex mixing every 10 min. Cells were centrifuged for 2.5 min at 16,100 × *g* at room temperature and washed twice with 1 mL of 37°C BHI broth, then resuspended in 600 µL BHI broth and incubated at 37°C in 5% $CO_2$ in the dark for 20 min. After incubation, cells were labeled with HADA (400 µM final), fixed, and prepared for vertical cell imaging as described above. Exposure times and %T settings were 5 ms and 40% for HADA and 5 ms and 50-100% for HT.

**Localization of bPBP2x with 2*R*,3*S*-β-lactone-L-Phe-fluorescein (7FL) in WT and Δ*pbp1b* mutant *Spn* cells.** WT (IU1945) and Δ*pbp1b* (E193) strains were labeled with 5 µg/mL of the bPBP2x-specific activity probe 7FL for 20 min in the dark as described in (7,8). Tilted, horizontally oriented cells were detected by 3D-SIM as described below.

**Localization of FDAA nodes (TADA labeling for 17 s) in vertically oriented *Spn* cells.** For experiments using short TADA pulse labeling to resolve FDAA nodes, cells from overnight cultures were diluted to $OD_{620}$ ≈0.02 in 2 mL of fresh BHI broth. At $OD_{620}$ ≈0.23, 600 µL cultures were labeled with TADA as described above, with the following changes: 250 µL of BHI broth containing TADA (45.5 µM final) was added to the filter and incubated for 3 s and filtered away, resulting in a total labeling time of ≈17 s. Cells were washed twice with 500 µL aliquots of room temperature PBS with filtering between washes. Cells were fixed and prepared for vertical cell imaging as described above. Exposure times and %T settings were 15 ms and 100% for TADA and 5 ms and 50% for sfGFP.

For strain IU8124 (Δ*ftsZ*//$P_{Zn}$-*ftsZ*$^+$), the following changes were made to the growth conditions: 0.2 mM $ZnCl_2$ (no $MnSO_4$) was added to overnight cultures. Following 12 h of incubation, serially diluted cultures of IU1945 and IU8124 still in exponential phase ($OD_{620}$ ≈0.05-0.4) were centrifuged at 16,100 × *g* for 5 m at room temperature, resuspended in 1 mL of 4°C PBS, and centrifuged again at room temperature. Pellets were resuspended in 2 mL of BHI broth, diluted to $OD_{620}$ ≈0.005 in 2 mL of BHI broth in a second tube (no

$ZnCl_2$), and incubated at 37°C in 5% $CO_2$. After a 2.5 h depletion of FtsZ, strains were labeled, fixed, and prepared for imaging as described above.

**Localization of TADA (17 s) followed by Boc-FL (21 s) in vertically oriented *Spn* cells.** For short pulses with TADA then Boc-FL, the following changes were made to the labeling procedure: Following the step in which the BHI broth containing TADA is filtered away, cells were washed with 500 µL of room temperature PBS and filtered. 250 µL of PBS containing 2 µg $mL^{-1}$ Boc-FL at 37°C was added to the filter, incubated for 7 s, and filtered away (total labeling time ≈21 s). Cells were washed with 500 µL of room temperature PBS, filtered, and resuspended in 600 µL of 4% (v/v) paraformaldehyde. Cells were fixed and prepared for vertical cell imaging as described above. Exposure time and %T setting was 15 ms and 100% for TADA and 6 ms and 50% for Boc-FL.

**Localization after sequential labeling with FDAAs BADA (green; 40 s), followed by TADA (red; 40 s), followed by HADA (blue; 40 s) in vertically oriented *Spn* cells.** For experiments with three tandem pulses of different colored FDAAs, cells from overnight cultures were diluted to $OD_{620}$ ≈0.02 in 2 mL of fresh BHI broth. At $OD_{620}$ ≈0.23, 600 µL of culture was centrifuged for 5 min at 16,100 × *g* at room temperature, and pellets were resuspended in 250 µL of BHI broth containing BADA (62.5 µM final). Cells were incubated for 40 s at 37°C, chilled for 20 s on dry ice, and centrifuged for 2.5 min at 16,100 × *g* at 4°C. After discarding the supernatant, the labeling, incubation, cooling, and centrifugation steps were repeated twice more with the same cells, first with TADA (62.5 µM final), and last with HADA (62.5 µM final). After discarding the supernatant from the HADA labeling, pellets were washed with 500 µL of ice-cold PBS, centrifuged, and resuspended in 600 µL of 4% (v/v) paraformaldehyde. Cells were fixed and prepared for vertical cell imaging as described above. Exposure time and %T settings were 5 ms and 50% for HADA, for BADA, and for TADA.

**Image acquisition by 3D-SIM.** A DeltaVision OMX Super Resolution system in the Indiana University Bloomington Light Microscopy Imaging Center (http://www.indiana.edu/~lmic/microscopes/index.html#OMX) was used for 3D-SIM. An Olympus PlanApo N 60X/1.42 oil objective was used, and images were acquired using a PCO.edge 4.2 (CMOS) camera system (Kelheim Germany). Laser lines were 405 nm with emission filters of 419–465 nm (for HADA), 488 nm with emission filters of 500–550 nm (for Boc-FL, sfGFP, and BADA), and 561 nm with emission filters of 609–654 nm (for Ceph-CT, TADA, and HT). For each sample tested, 3D-SIM of horizontally oriented cells was performed to ensure that the refractive index of the immersion oil used matched the refractive index of the sample prepared (1.518 oil for samples prepared in hardset antifade and 1.514 oil for samples prepared with Slowfade gold antifade reagent). To initially focus cells, the 561 nm laser line was used with the lowest laser power and exposure times needed to detect TADA-labeled rings in regions of interest. To acquire images for 3D-SIM, between 9-15 *z* sections (each 0.125 µm thick) were initially imaged using the laser powers indicated above. Image processing was performed with DeltaVision SoftWoRx Software (GE Healthcare), consisting of OMX Reconstruction (Wiener filter value set at 0.001) and OMX Align. Vertically oriented *Spn* cells were caught at different stages of the cell cycle. To eliminate TADA and other labeling outside of midcell, all final reconstructed images in this work are cross sections of 2-4 z sections (0.25-0.50 µm total thickness) containing only division planes, unless noted otherwise.

**Analysis of FDAA node distribution using a custom Matlab GUI.** Reconstructed 3D-SIM image slices 2-4 *z* sections thick were analyzed using a point-and-click vertical image analysis graphical user interface (VIMA-GUI) organized in MATLAB (The Mathworks). Briefly, several arbitrary points were selected around the division site of a single image, and these points were used to create a marker circle around the division site to determine pixel intensities. Nodes in each image were then manually selected, and these designations were used to reformat the circle into a size and position that overlaid the division site as closely as possible for each individual image. Based on the location of the marked nodes, the program placed a pair of concentric circles with a 5-pixel gap between them, such that the fluorescent division site ring was aligned between the concentric rings. The program then measured the intensity and distribution of fluorescence within the space between the concentric rings. Nodes were selected as individual foci, which could be oblong or round shaped. Oblong foci that demonstrated no partial segmentation were considered as one focus. Oblong patterns that contained distinct multiple foci, even if slightly connected, were counted as separate foci. A subroutine in the VIMA-GUI program was used to calculate the correlation coefficients of the number of nodes and their spatial distribution in dual-labeled cells.

**Widefield fluorescence microscopy.** A DeltaVision personalDV microscope (Applied Precision) equipped with a CoolSNAP HQ2/HQ2-ICX285 camera was used to perform widefield fluorescence microscopy. An 8 s exposure was taken with 100% T for imaging TADA labeling. Samples were prepared as described above for 17 s pulse labeling with TADA and visualization of vertically oriented cells. Where indicated, image deconvolution processing was performed with DeltaVision SoftWoRx Software, including adjustments to Wiener filter settings.

**Expression and purification of *Spn* bPBP2x and bPBP2b soluble fragments for polyclonal antibody production.** *Plasmid construction.* A DNA fragment of *pbp2b* (*spd_1486*) encoding amino acids Q40 to N685 of the extracellular domain of bPBP2b was amplified from D39W Δ*cps* genomic DNA using primers: oAT90 (GTACATATGCAGGTTTTGAACAAGGATTTTTACG) and oAT91 (ACTAAGCTTATTCATTGGATGGTATTTTTGATACAGA), where restriction sites are underlined. After digestion with *Nde*I and *Hind*III, the PCR amplicon product was ligated into vector pET22b, resulting in plasmid pATPL005 that imparts resistance to ampicillin (Amp$^{R}$) and expresses *Spn* PBP2b$^{Q40\text{-}N685}$-His$_{6}$. Similarly, a DNA fragment of *pbp2x* (*spd_0306*) encoding amino acids G49 to D750 of the extracellular domain of bPBP2x was amplified from D39W Δ*cps* genomic DNA using primers: oAT92 (GTACATATGGGGACAGGCACTCGCTTTG) and oAT93 (ACTAAGCTTGTCTCCTAAAGTTAATGTAATTTTTTTAATG), where restriction sites are underlined. After digestion with *Nde*I and *Hind*III, the PCR amplicon product was ligated into vector pET22b, resulting in plasmid pATPL025 that imparts Amp$^{R}$ and expresses *Spn* PBP2x$^{G49\text{-}D750}$-His$_{6}$.

*Expression, protein purification, and antibody production*. Each plasmid was transformed into *E. coli* BL21(DE3), and the resulting strains were grown in 1 L of LB broth supplemented with 50 mg/L carbenicillin at 37ºC with shaking to $OD_{600}$ ≈0.4. Culture were cooled to 16ºC before inducing protein expression with 500 μM IPTG. Cells were harvested 18 h post-induction by centrifugation (4,200 *x g* for 15 min at 4ºC). Pelleted cells were resuspended in 30 mL lysis buffer (50 mM HEPES pH 7.5, 500 mM NaCl) supplemented with 1 mM PMSF, and resuspended cells were lysed by three passages through a cell disruptor (Emulsiflex C5, Avestin) at 15,000 psi. Insoluble material was removed by ultracentrifugation (100,000 *x g* for 30 min at 4ºC). 1 mL pre-equilibrated Ni-NTA resin (Qiagen) was added to each supernatant, and mixtures were stirred for 1 h at 4ºC. Sample were loaded onto a gravity column and washed with 20 mL wash buffer (500 mM HEPES pH 7.5, 500 mM NaCl, 0.1% (vol/vol) Triton X-100) containing 20 mM imidazole, and then 20 mL wash buffer containing 50 mM imidazole. Proteins were eluted in 5 mL wash buffer containing 300 mM imidazole. The eluate was further purified by size exclusion chromatography with a Superdex 200 10/300 GL column (GE Healthcare) equilibrated in running buffer (50 mM HEPES pH 7.5, 150 mM NaCl, 0.1% (vol/vol) Triton X-100-Reduced). Fractions containing the target proteins were concentrated by centrifugal filtration and protein concentrations were determined with Pierce BCA Protein Assay Kit (Thermo Fisher Scientific). The purity of soluble *Spn* bPBP2b and bPBP2x fragments was >95% based on Coomassie Blue staining of PAGE gels. Purified PBP fragments were used to generate rabbit polyclonal antibodies (Covance). For this study, anti-bPBP2x and anti-bPBP2b were used without further purification. Western blotting showed that both antibody preparations showed little non-specific background, and comparisons of blots of native PBPs with PBP-fusion constructs ruled out the presence of overlapping non-specific bands at the molecular mass of the PBP or the PBP-fusion constructs (Fig. S3C).

### SI APPENDIX, ADDITIONAL DATA

**Labeling patterns of fluorescent-cephalosporin (Ceph-CT) and fluorescent-bocillin (Boc-FL) overlap in wild-type *Spn* cells.** A previous study reported the synthesis and characterization of new fluorescent β-lactam probes for PBPs in *Bacillus subtilis* (*Bsu*) and *Spn* (9). Cells of an unencapsulated (Δ*cps*) derivative (IU1945) of progenitor serotype 2 strain D39 were labeled sequentially with cephalosporin conjugated to TAMRA (Ceph-CT), followed by fluorescent-bocillin (Boc-FL) for relatively long times and viewed by 2D-EFM or 3D-SIM (9). Ceph-CT preferentially labels aPBP2a, aPBP1b, and class C cPBP3, whereas Boc-FL labels all six *Spn* PBPs nonspecifically (9,10). Limited 3D-SIM imaging showed incomplete, nonoverlapping labeling by Ceph-CT and Boc-FL at the midcell septa and equatorial rings of dividing *Spn* cells (9). Although this study established the utility of Ceph-CT and related β-lactam probes, the small number of foci labeled by nonspecific Boc-FL seemed counterintuitive.

We reprised these experiments by optimizing labeling conditions and exposure settings to minimize photobleaching during 3D-SIM (see method below). Contrary to the previous result, labeling cells sequentially with Ceph-CT (30 min) followed by Boc-FL (10 min) gave contiguous labeling of each probe with substantial overlap around the midcell of predivisional *Spn* cells (Fig. S6). Gaps, blurring, and non-uniformity (“horse collars”) in the lower 3D-SIM images are indicative of lower axial resolution during rotation (see *Introduction*). We conclude that over the long labeling times used in this experiment, active PBPs distribute over the entire equatorial ring of predivisional *Spn* cells and are not located in a small number of nonoverlapping puncta.

*Method of saturating labeling of Spn cells with Boc-FL and Ceph-CT.* Saturating labeling with Boc-FL and Ceph-CT was performed as described in (9) with the following changes. Briefly, WT cells from overnight cultures were diluted to $OD_{620}$ ≈0.02 in 2 mL fresh BHI broth and incubated as described above. At $OD_{620}$ ≈ 0.23, 500 μL of culture was centrifuged at 16,100 × *g* for 5 min at room temperature. The pellet was resuspended in 25 μL of 1X PBS (PBS; Ambion, AM9625) containing Ceph-CT (10 μg $mL^{-1}$ final) for 30 min at room temperature, centrifuged at 16,100 × *g* for 5 min at room temperature, and resuspended in 25 μL of PBS containing Boc-FL (5 μg $mL^{-1}$ final) for 10 min at room temperature. Cells were centrifuged at 16,100 × *g* for 5 m at room temperature, washed with 500 μL of ice-cold PBS, centrifuged again, and fixed by resuspending in 500 μL of 4% (v/v) paraformaldehyde (EMS; 157-4) and incubated in the dark for 15 min at room temperature and 45 min on ice. Fixed cells were centrifuged (16,100 × *g* for 5 min at room temperature), washed with 500 μL of ice-cold PBS, centrifuged again, and resuspended in 50 μL ice-cold GTE buffer (50 mM glucose, 1 mM EDTA, 20 mM Tris-HCl, pH 7.5). The entire cell suspension was deposited on a coverslip, incubated for 5 min in the dark at room temperature, and aspirated via a vacuum pump. The cells on the coverslip were washed for 10 s with 70 μL PBS-T (0.2% (v/v) Triton X-100 (Mallinckrodt, H282) in PBS) and aspirated. The coverslip was air dried, immersed in -20°C methanol for 10 min, removed, and air dried. The coverslip was incubated with 70 μL PBS-T for 5 min, aspirated, and briefly washed twice with 70 μL PBS. The coverslip was allowed to air dry and 7 μL of Slowfade gold antifade reagent (Invitrogen, S36936) was applied to the coverslip, which was then inverted onto a microscope slide (VWR, 16004-368) and sealed with clear nail polish. Image acquisition was performed as described above, with the following exposure times and %T: 5 ms and 50% for Boc-FL and for Ceph-CT.

### SI APPENDIX, TABLES

**Table S1.** *Streptococcus pneumoniae* strains used in this study

| Strain | Genotype (description and linkers)[a] | Antibiotic resistance[b] | Reference or source |
|---|---|---|---|
| E193 | D39 Δ*cps* Δ*pbp1b*:: $P_c$-*erm* | $Erm^R$ | (2) |
| E751 | D39 Δ*cps* Δ*khpA*::$P_c$-*erm* | $Erm^R$ | (11) |
| EzrA-Ts1[c] | D39 Δ*cps* *ezrA*(T506I)-$P_c$-*erm* (IU1945 X Error Prone PCR product) | $Erm^R$ | This study |
| K164 | D39 Δ*cps* Δ*pbp1a*::$P_c$-[*kan-rpsL*$^+$] | $Kan^R$ | (12) |
| IU1690 | D39W *cps*$^+$ | None | (13,14) |
| IU1824 | D39W Δ*cps* *rpsL1* | $Str^R$ | (13) |
| IU1945 | D39W Δ*cps* | None | (13) |
| IU5456 | D39 Δ*cps* *ezrA*-$L_0$-FLAG$^3$-$P_c$-*erm* | $Erm^R$ | (15) |
| IU5838 | D39 Δ*cps* *gpsB*-FLAG-$P_c$-*erm* | $Erm^R$ | (2) |
| IU6545 | D39 Δ*cps* *ezrA*-HA-$P_c$-*erm* (IU1945 X fusion amplicon *ezrA*-HA-$P_c$-*erm)* | $Erm^R$ | This study |
| IU8124[d] | D39 Δ*cps* Δ*ftsZ*::*aad9* //Δ*bgaA*::*tet*-t1t2-$P_{Zn}$-*ftsZ*$^+$ | $Spc^R$ $Tet^R$ | (16) |
| IU8872 | D39 Δ*cps* Δ*bgaA*::*tet*-$P_{Zn}$-RBS$^{mltG}$-*mltG*$^+$ | $Tet^R$ | (17) |
| IU9023 | D39 Δ*cps* *rpsL1* $P_c$-[*kan-rpsL*$^+$]-*pbp2b*$^+$ | $Kan^R$ | (16) |
| IU9621 | D39 Δ*cps* *rpsL1* Δ*khpA*//Δ*bgaA*::*kan*-t1t2-$P_{ftsA}$-*khpA*$^+$ | $Str^R$ $Kan^R$ | (11) |
| IU9689 | D39 Δ*cps* Δ*bgaA*::*kan*-t1t2-$P_{Zn}$-*khpA*$^+$ (IU1945 X amplicon Δ*bgaA*::*kan*-t1t2-$P_{Zn}$-*khpA*$^+$) | $Kan^R$ | This study |
| IU9783 | D39 Δ*cps* *rpsL1* *mltG*(Y488D) Δ*pbp2b*<>*aad9* | $Str^R$ $Spc^R$ | (17) |
| IU9805 | D39 Δ*cps* Δ*bgaA*::*kan*-t1t2-$P_{Zn}$-*sepF*$^+$ (IU1945 X fusion amplicon Δ*bgaA*::*kan*-t1t2-$P_{Zn}$-*sepF*$^+$) | $Kan^R$ | This study |
| IU9965 | D39 Δ*cps* *rpsL1* *sfgfp*-$L_1$-*pbp2b* markerless | $Str^R$ | (16) |
| IU9985 | D39 Δ*cps* *rpsL1* *ftsZ*-$L_2$-*sfgfp* markerless | $Str^R$ | (16) |
| IU10447 | D39 Δ*cps* *ezrA*$^+$-$P_c$-*erm* (IU1945 X fusion amplicon *ezrA*$^+$-$P_c$-*erm*) | $Erm^R$ | This study |
| IU10612 | D39 Δ*cps* *ftsZ*(G107S) markerless //Δ*bgaA*::*tet*-$P_{Zn}$-*ftsZ*$^+$ | $Str^R$ | (16) |
| IU11034 | D39 Δ*cps* *ezrA*(T506I)-HA-$P_c$-*erm* (IU1945 X fusion amplicon ezrA(T506I)-HA-$P_c$-*erm*) | $Erm^R$ | This study |
| IU11051 | D39 Δ*cps* *gpsB*$^+$-$P_c$-*erm* | $Erm^R$ | This study |

| | | | |
|---|---|---|---|
| | (IU1945 X fusion amplicon *gpsB*$^+$-$P_c$-*erm*) | | |
| IU11155[c] | D39 Δ*cps gpsB*(K96N)-$P_c$-*erm*<br>(IU1945 X Error Prone PCR product) | Erm$^R$ | This study |
| IU11157 | D39 Δ*cps rpsL1 isfgfp*-$L_1$-*pbp2x* markerless | Str$^R$ | (16) |
| IU11303 | *Streptococcus mitis* ATCC 49456<br>Strain NCTC 12261 [NS 51] | Not indicated | (18) |
| IU11956 | D39 Δ*cps gpsB* (K96N)-$P_c$-*erm* (IU1945 X amplicon IU11155) | Erm$^R$ | This study |
| IU12376[d] | D39 Δ*cps rpsL1* Δ*ftsX::*$P_c$-*aad9*<br>//Δ*bgaA*::*tet*-t1t2-$P_{Zn}$-*ftsX*$^+$ | Spec$^R$ Str$^R$ Tet$^R$ | (6) |
| IU12575[d] | D39 Δ*cps rpsL1* Δ*ftsX::*$P_c$-[*kan-rpsL*$^+$]<br>//Δ*bgaA*::*tet*-$P_{Zn}$-*ftsX*$^+$ (IU12376 X fusion amplicon Δ*ftsX*::$P_c$-[*kan-rpsL*$^+$]) | Tet$^R$ Kan$^R$ | This study |
| IU13333 | *Enterococcus faecalis* Strain NJ-3,<br>ATCC 51299 | Van$^R$ Tei$^S$ | ATCC |
| IU14738 | D39 Δ*cps rpsL1 iht*-$L_6$-*mapZ markerless* | Str$^R$ | (16) |
| IU14927 | D39 Δ*cps rpsL1 iht*-$L_6$-*pbp2*x markerless | Str$^R$ | (16) |
| IU15364 | D39 Δ*cps rpsL1 ftsX*$_{(1\text{-}77)}$-*iht*-$L_6$-*ftsX*$_{(80\text{-}308aa)}$<br>//Δ*bgaA*::*tet*-t1t2-$P_{Zn}$-*ftsX*$^+$ markerless<br>(IU12575 X fusion amplicon *ftsX*$_{(1\text{-}77aa)}$-*iht*-$L_6$-*ftsX*$_{(80\text{-}308aa)}$) | Str$^R$ Tet$^R$ | This study |
| IU15928 | D39 Δ*cps rpsL1 iht*-$L_6$-*pbp2b* markerless<br>(IU9023 X fusion amplicon *iht*-$L_6$-*pbp2b)* | Str$^R$ | This study |
| IU16121[e] | D39 Δ*cps rpsL1*<br>*ftsX*$_{(1\text{-}77aa)}$-*isfgfp*-$L_6$-*ftsX*$_{(80\text{-}308aa)}$ markerless<br>//Δ*bgaA*::*tet*-t1t2-$P_{Zn}$-*ftsX*$^+$<br>(IU12575 transformed X fusion amplicon *ftsX*$_{(1\text{-}77aa)}$-*isfgfp*-$L_6$-*ftsX*$_{(80\text{-}308aa)}$) | Str$^R$ Tet$^R$ | This study |
| IU16499 | D39 Δ*cps rpsL1*<br>Δ*bgaA*::*kan*-t1t2-$P_{Zn}$-*iht-pbp2b*<br>(IU1824 X fusion amplicon *ΔbgaA*::*kan*-t1t2-$P_{Zn}$-*iht*-$L_6$-*pbp2b*) | Kan$^R$ | This study |
| IU16553 | D39 Δ*cps rpsL1 iht*-$L_6$-*pbp2b* markerless *//*<br>*ΔbgaA*::*kan*-t1t2-$P_{Zn}$-*iht*-$L_6$-pbp2b<br>(IU15928 X amplicon *ΔbgaA*::*kan*-t1t2-$P_{Zn}$-*iht*-$L_6$-*pbp2b* from IU16499) | Str$^R$ Kan$^R$ | This study |

[a]Methods to construct strains are described in (19,20). L refers to linkers used with the following amino acid sequences: $L_0$:(GSAGSAAGSG), $L_1$:(LEGSG), $L_2$:(KLDIEFLQ), and $L_6$ :(LEGSGQGPGSGQGSG). *i* in *isfgfp* or *iht* refers to an i-tag sequence, used to increase protein expression as described in (21). *ht* refers to HaloTag protein with a *Spn*-optimized coding sequence (16). The amino acid sequence of the FLAG tag is

DYKDDDDK, described in (5,22). The amino acid sequence of the HA tag is YPYDVPDYA (2,23). The $P_c$-[*kan-rpsL*$^+$] Janus cassette used for allelic exchange is reported in (24).

[b]Antibiotic resistance markers: Erm$^R$; erythromycin, Kan$^R$; kanamycin, Spc$^R$; spectinomycin, Str$^R$; streptomycin, Tet$^R$; tetracycline, Amp$^R$; ampicillin, and Van$^R$; vancomycin. Antibiotic sensitive: Tei$^S$; teicoplanin.

[c]Error-prone PCR mutagenesis was performed starting with amplification of the *ezrA*$^+$-$P_c$-*erm* or *gpsB*$^+$-$P_c$-*erm* region of strain IU10447 or IU11051 to obtain *ezrA*-Ts1 (*ezrA*(T506I)-$P_c$-*erm*) or IU11155 (*gpsB*(K96N)-$P_c$-*erm*), respectively. Error-prone PCR mutagenesis was performed using the GeneMorph II Random Mutagenesis Kit (C# 200550) per the manufacturer's instructions to obtain a low mutation frequency of 0-4.5 mutations/kb. Resulting mutations-containing amplicons (amplicons*) were transformed into IU1945 as previously described in (19). Transformants were plated onto TSAII-BA plates containing erythromycin at 32°C, and colonies were screened for temperature sensitivity at 42°C on the same kind of plate. Isolates of EzrA-Ts1 and IU11155 grew readily at 32 °C, but were unable to grow at 42° when patched onto TSAII-BA or inoculated into BHI broth at 42°C. Sequencing revealed that EzrA-Ts1 contains a point mutation: (ACA→ATA) resulting in *ezrA*(T506I) and IU11155 contains a point mutation: (AAA→AAC) resulting in *gpsB*(K96N). Backcross experiments were performed using purified and sequenced amplicons*, and resulting strains confirmed temperature sensitivity at 42°C on TSAII-BA plates and in BHI broth. An HA tag was added to e*zrA*(T506I) to give strain IU11034 (*ezrA*(T506I)-HA-$P_c$-*erm*) used in experiments.

[d]To eliminate selective pressure, depletion strains were supplemented with inducer throughout all stages of construction and storage. The required concentrations of inducer for specific isolates are as follows: IU8124 requires added 0.3 mM $ZnCl_2$ and 0.03 mM $MnSO_4$ to induce ectopic expression of *ftsZ*$^+$, although complementation still occurs using 0.2 mM $ZnCl_2$ alone; IU12376 and IU12575 require added 0.45 mM $ZnCl_2$ and 0.045 mM $MnSO_4$ to induce ectopic *ftsX*$^+$.

[e] Strain IU16121 (*ftsX'-isfgfp-ftsX'* // $P_{Zn}$-*ftsX*$^+$) did not require zinc for growth, and zinc and manganese were not added to media for this strain.

**Table S2**. Oligonucleotide primers used for construction of *Spn* strain in this study

| Primer | Sequence (5' to 3') | Template[a] | Amplicon Product |
|---|---|---|---|
| **For construction of EzrA-Ts1 (*ezrA*(T506I)-$P_c$-*erm*)** | | | |
| AL295 | CCCAAATCCACAGTTTGAAGGACAAACG | IU10447 | Mixed mutation pool, error prone PCR mutagenesis amplicons[c] |
| AL297 | GGACCTACTCCTATTGGAGCCCAAC | | |
| **For construction of IU6545 (*ezrA*-HA-$P_c$-*erm*)** | | | |
| TT192 | ATCGTGTTCCAGCCTTGGTTACGACGCT TT | IU1690 hence referred to as D39 | 5' fragment containing *ezrA*-HA |
| SV005 | CCCGGTTAAGCATAATCTGGAACATCAT ATGGATAAAAACGAATCGTTTCACGTGTT TTC | | |
| SV006 | GATTCGTTTTTATCCATATGATGTTCCAG ATTATGCTTAACCGGGCCCAAAATTTGTT TG | IU5456 | 3' fragment containing HA-$P_c$-*erm* and downstream of *ezrA* |
| AL297 | GGACCTACTCCTATTGGAGCCCAAC | | |
| **For construction of IU9689 (Δ*bgaA*::*kan*-t1t2-$P_{Zn}$-*khpA*$^+$) (an intermediate)** | | | |
| P146 | TGGCCATTCATCGCTGGTCGTGCTGAAA T | IU9621 | 5' fragment containing Δ*bgaA*::*kan*-t1t2-$P_{Zn}$ |
| JQ145 | CCGTATCAGCAAAACCAAAAAAGCCATC TAGTAGAAACGCAAAAAGGCCATCCGTC AGGA | | |

| | | | |
|---|---|---|---|
| JQ146 | TCCTGACGGATGGCCTTTTTGCGTTTCTACTAGATGGCTTTTTTGGTTTTGCTGATACGG | IU8872 | $RBS_{mltG}$ |
| JQ147 | CGCAATAATGAGATTTTCAATCGTATCCATAAGTTTTTCCTCCTTGTTGATAATTTGTTA | | |
| JQ148 | TAACAAATTATCAACAAGGAGGAAAAACTTATGGATACGATTGAAAATCTCATTATTGCG | IU9621 | 3' fragment end of *khpA-bga'* to downstream |
| CS121 | GCTTTCTTGAGGCAATTCACTTGGTGC | | |
| **For construction of IU9805 (Δ*bgaA*::*kan*-t1t2-$P_{Zn}$-*sepF*$^+$) (an intermediate)** | | | |
| P146 | TGGCCATTCATCGCTGGTCGTGCTGAAAT | IU9689 | 5' fragment containing Δ*bgaA*::*kan*-t1t2-$P_{Zn}$-$RBS_{ftsA-}$ |
| AJP32 | ACATCGCTTCCTCTCTATCTTCCTTGTTATAATAGATTTATGAACACCTTGTTCATTATC | | |
| AJP107 | GGAAGATAGAGAGGAAGCGATGTAATGTCTTTAAAAGATAGATTCGATAGATTTATAGAT | D39 | *sepF*+ |
| AJP108 | CAACTGGTTTATGAGAAAGTAAGTTCTTTTATCGTACTCTATTTCGCTTCATATCAAAA | | |
| AJP109 | GATATGAAGCGAAATAGAGTACGATAAAAGAACTTACTTTCTCATAAACCAGTTGCTG | D39 | 3' fragment end of *sepF-bga'* to downstream |
| CS121 | GCTTTCTTGAGGCAATTCACTTGGTGC | | |
| **For construction of IU10447 (*ezrA*$^+$-$P_c$-*erm*)** | | | |
| TT192 | ATCGTGTTCCAGCCTTGGTTACGACGCTTT | D39 | 5' fragment containing only 3' of *ezrA*$^+$ |
| AJP134 | AACAAATTTTGGGCCCGGTTAAAAACGAATCGTTTCACGTGTTTTCT | | |
| AJP135 | AACACGTGAAACGATTCGTTTTTAACCGGGCCCAAAATTTGTTTGATTT | IU6545 | 3' fragment containing $P_c$-*erm* and downstream of *ezrA* |
| TT330 | TTCTTGAAGGAGAGAGTCGAGTCCGAACTCCTC | | |
| **For construction of IU11034 (*ezrA*(T506I)-HA-$P_c$-*erm*)** | | | |
| AL295 | CCCAAATCCACAGTTTGAAGGACAAACG | EzrA-Ts1 | 5' fragment containing *ezrA*(T506I)-HA |
| SV005 | CCCGGTTAAGCATAATCTGGAACATCATATGGATAAAAACGAATCGTTTCACGTGTTTTC | | |
| SV006 | GATTCGTTTTTATCCATATGATGTTCCAGATTATGCTTAACCGGGCCCAAAATTTGTTTG | IU6545 | 3' fragment containing HA-$P_c$-*erm* and downstream of *ezrA* |
| TT330 | TTCTTGAAGGAGAGAGTCGAGTCCGAACTCCTC | | |

| | | | |
|---|---|---|---|
| **For construction of IU11051 (*gpsB*$^{+}$-$P_c$-*erm*)** | | | |
| TT196 | GCCAAGCCCTGAGACAAATAGTAGTCGTTGGT | D39 | 5’ fragment containing *gpsB*$^{+}$ |
| TT905 | ACAAATTTTGGGCCCGGTTAAAAATCTGAGTTATCTAAAATTTGTTTACCAAA | | |
| TT906 | GTAAACAAATTTTAGATAACTCAGATTTTTAACCGGGCCCAAAATTTGTTTGAT | IU5838 | 3’ fragment containing $P_c$-*erm* and downstream of *gpsB*$^{+}$ |
| TT197 | TTTGATACGATCTGCTGCCCGAAGCCAAAGGT | | |
| **For construction of IU11155 (*gpsB*(K96N)-$P_c$-*erm*)** | | | |
| TT196 | GCCAAGCCCTGAGACAAATAGTAGTCGTTGGT | IU11051 | Mixed mutation pool, error prone PCR mutagenesis amplicons[c] |
| TT197 | TTTGATACGATCTGCTGCCCGAAGCCAAAGGT | | |
| **For construction of IU12575 (Δ*ftsX*::$P_c$-[*kan-rpsL*$^{+}$])** | | | |
| KB250 | ATTGTTGTCCAATCAACAGTGGATCGTACC | D39 | 5’ fragment containing 60 bp of *ftsX* |
| BR176 | CACATTATCCATTAAAAATCAAACGGATCCTAACCATTTCGTTTCAAACTTTTTAAGGCT | | |
| KanrpsL forward | TAGGATCCGTTTGATTTTTAATGGATAATG | $P_c$-[*kan-rpsL*$^{+}$] cassette | Middle fragment containing $P_c$-[*kan-rpsL*$^{+}$] cassette |
| KanrpsL reverse | GGGCCCCTTTCCTTATGCTTTTG | | |
| BR177 | CGGTACTAAACGTCCAAAAGCATAAGGAAAGGGGCCCGGGGTTTTCATTGGTTCATTGGG | D39 | 3’ fragment containing 60 bp of *ftsX* |
| KB251 | GCGATAACTACCATCAAAGCCTTACCGAAT | | |
| **For construction of IU15364 (*ftsX*$_{(1\text{-}77aa)}$-*iht*-$L_6$-*ftsX*$_{(80\text{-}308aa)}$)** | | | |
| KB250 | ATTGTTGTCCAATCAACAGTGGATCGTACC | D39 | 5’ fragment containing upstream of *ftsX* and 5’ of *ftsX* up to 77 codons and *itag* |
| BR279 | CCAAAAAAATTTCCAAACCTTTTTTATCTTTTTCAATTGTCTGACTATTATCTTCCACAT | | |
| BR278 | GTGGAAGATAATAGTCAGACAATTGAAAAAGATAAAAAAGGTTTGGAAATTTTTTTGGCT | IU14738 | Middle fragment containing *iht*-$L_6$ |
| BR281 | CTTGTGGTAGTCATTATTTGTAACAGTTTGACCAGAACCTTGACCAGATCCTGGTCCTTG | | |

| | | | |
|---|---|---|---|
| BR280 | AAGGACCAGGATCTGGTCAAGGTTCTGGTCAAACTGTTACAAATAATGACTACCACAAGG | D39 | 3’ fragment containing 3’ of *ftsX* (codons 80-308) and downstream of *ftsX* |
| KB251 | GCGATAACTACCATCAAAGCCTTACCGAAT | | |
| **For construction of IU15928 (*iht*-$L_6$-*pbp2b* markerless)** | | | |
| TT452 | GGAGGGTTGGCTGTGGGTGGCTACAAGAAC | D39 | 5’ frag upstream of *pbp2b* |
| ML82 | AGCCAAAAAAATTTCCAAACCTTTTTATCCATTTCTAACTTAAAATCTTACTCTTAATT | | |
| AJP405 | GATAAAAAAGGTTTGGAAATTTTTTTGGCTTCTGCTGAAATTGGTACTGGTTTTCCATTT | IU14738 | Middle fragment containing *iht*-$L_6$ |
| YT104 | ACCAGAACCTTGACCAGATCCTGGTCCTTG | | |
| ML83 | CAAGGACCAGGATCTGGTCAAGGTTCTGGTAGACTGATTTGTATGAGAAAATTTAACAGC | D39 | 3’ frag *pbp2b* |
| TT352 | TGAAGGACTGGAAAGACCACTGCACCTTCT | | |
| **For construction of IU16121 (*ftsX*$_{(1\text{-}77aa)}$-*isfgfp*-$L_6$-*ftsX*$_{(80\text{-}308aa)}$)** | | | |
| KB250 | ATTGTTGTCCAATCAACAGTGGATCGTACC | IU15364 | 5’ fragment upstream and 5’ of *ftsX* up to codon 77 and *itag* |
| KB540 | ACCTGTGAACAGCTCTTCTCCTTTCATAGAAGCCAAAAAAATTTCCAAACCTTTTTATC | | |
| TT900 | ATGAAAGGAGAAGAGCTGTTCACAGGTGTTGTGCCGAT | IU9985 | Middle fragment containing *sfgfp* |
| KB541 | ATCCTGGTCCTTGTCCTGATCCTTCCAACTTATAAAGCTCATCCATGCCGTGAGTGATAC | | |
| KB542 | TTGGAAGGATCAGGACAAGGACCAGG | IU15364 | 3’ fragment 3’ of *ftsX*-$L_6$-codons 80-308aa and downstream of *ftsX* |
| KB251 | GCGATAACTACCATCAAAGCCTTACCGAAT | | |
| **For construction of IU16499 (Δ*bgaA*::*kan*-t1t2-$P_{Zn}$-*iht*-$L_6$-*pbp2b*)** | | | |
| P146 | TGGCCATTCATCGCTGGTCGTGCTGAAAT | IU9805 | 5’ fragment upstream to Δ*bga*::*kan*-t1t2-$P_{Zn}$ |
| TT1260 | AAATTTCCAAACCTTTTTATCCATTACATCGCTTCCTCTCTATCTTCCTTGTTA | | |

| | | | |
|---|---|---|---|
| TT1261 | GGAAGATAGAGAGGAAGCGATGTAATG GATAAAAAAGGTTTGGAAATTTTTTTG | IU15928 | *iht*-$L_6$-*pbp2b* |
| BR72 | AACTGGTTTATGAGAAAGTAAGTTCTTCT AATTCATTGGATGGTATTTTTGATACAGA TT | | |
| BR71 | GTATCAAAAATACCATCCAATGAATTAGA AGAACTTACTTTCTCATAAACCAGTTGCT GC | D39 | 3’ fragment *bgaA’* to downstream |
| CS121 | GCTTTCTTGAGGCAATTCACTTGGTGC | | |

**Table S3.** Compilation (mean ± SD) of ring diameters, nodes per ring, and arc distances between nodes for this study

| | **Strain genotype and/or relevant condition**[a] [b] | **Ring diameter (μm) (# of cells analyzed )** | **Nodes per ring (# of cells analyzed)** | **Arc distance between nodes (μm) (# of arcs analyzed)** |
|---|---|---|---|---|
| 1 | *isfgfp-pbp2x*<br>Fig. 2 (sfGFP nodes) | 0.80 ± 0.07<br>(29) | 11.97 ± 1.97<br>(29) | 0.21 ± 0.08<br>(347) |
| 2 | *sfgfp-pbp2b*<br>Fig. 2 (sfGFP nodes) | 0.88 ± 0.09<br>(29) | 10.90 ± 1.68<br>(29) | 0.25 ± 0.11<br>(316) |
| 3 | **Wild-type (WT)**<br>Fig. 3 | 0.84 ± 0.05<br>(76) | 9.82 ± 1.03<br>(76) | 0.27 ± 0.07<br>(746) |
| 4 | FtsZ depleted-Reg.<br>Fig. 5 | 1.18 ± 0.12<br>(19) | 12.11 ± 2.28<br>(19) | 0.31 ± 0.11<br>(230) |
| 5 | FtsZ depleted-Irreg.<br>Fig. 5 | 1.15 ± 0.17<br>(28) | 10.46 ± 2.13<br>(28) | 0.35 ± 0.22<br>(293) |
| 6 | Δ*pbp1a*<br>Fig. 6, Fig. S4 | 0.64 ± 0.08<br>(52) | 7.21 ± 1.02<br>(52) | 0.28 ± 0.09<br>(376) |
| 7 | Δ*khpA*<br>Fig. 6, Fig. S4 | 0.71 ± 0.09<br>(54) | 8.39 ± 1.20<br>(54) | 0.27 ± 0.08<br>(453) |
| 8 | *ftsZ*(G107S)-Reg.<br>Fig. 6, Fig. S4 | 0.96 ± 0.12<br>(24) | 11.17 ± 1.58<br>(24) | 0.27 ± 0.09<br>(268) |
| 9 | *ftsZ*(G107S)-Irreg.<br>Fig. 6, Fig. S4 | 0.99 ± 0.10<br>(46) | 10.11 ± 1.37<br>(46) | 0.31 ± 0.12<br>(465) |
| 10 | *ezrA*(T506I)-Reg.<br>Fig. 6, Fig. S4 | 0.99 ± 0.14<br>(29) | 11.03 ± 1.72<br>(29) | 0.28 ± 0.09<br>(320) |
| 11 | *ezrA*(T506I)-Irreg.<br>Fig. 6, Fig. S4 | 1.02 ± 0.16<br>(30) | 10.23 ± 1.89<br>(30) | 0.31 ± 0.13<br>(307) |
| 12 | *gpsB*(K96N)-Reg.<br>Fig. 6, Fig. S4 | 1.12 ± 0.07<br>(20) | 12.30 ± 1.90<br>(20) | 0.28 ± 0.09<br>(246) |
| 13 | *gpsB*(K96N)-Irreg.<br>Fig. 6, Fig. S4 | 1.11 ± 0.10<br>(34) | 10.29 ± 1.57<br>(34) | 0.34 ± 0.15<br>(350) |
| 14 | *sfgfp-pbp2b*<br>Fig. 7 | 0.86 ± 0.08<br>(48) | 9.35 ± 1.06<br>(48) | 0.29 ± 0.09<br>(449) |
| 15 | *sfgfp-pbp2b*<br>Fig. 7 (sfGFP nodes) | 0.84 ± 0.07<br>(48) | 9.19 ± 1.38<br>(48) | 0.29 ± 0.11<br>(441) |
| 16 | WT + Boc-FL<br>Fig. 7 | 0.83 ± 0.06<br>(18) | 9.39 ± 1.09<br>(18) | 0.28 ± 0.08<br>(169) |
| 17 | WT + Boc-FL<br>Fig. 7 (Boc-FL nodes) | 0.85 ± 0.07<br>(18) | 10.61 ± 1.20<br>(18) | 0.25 ± 0.08<br>(191) |
| 18 | *S. mitis*<br>Fig. 8 | 0.75 ± 0.04<br>(19) | 9.05 ± 1.03<br>(19) | 0.26 ± 0.08<br>(172) |
| 19 | *E. faecalis*<br>Fig. 8 | 0.77 ± 0.04<br>(20) | 8.90 ± 0.72<br>(20) | 0.27 ± 0.07<br>(178) |

[a]TADA nodes were quantified unless noted otherwise.

[b]Strains used: K164 (Δ*pbp1a*), E751(Δ*khpA*), IU1945 (WT), IU10612 (*ftsZ*(G107S)), IU11034 (*ezrA*(T506I)), IU11956 (*gpsB*(K96N)), IU8124 (FtsZ depleted), IU11157 (*isfgfp-pbp2x*), IU9965 (*sfgfp-pbp2b*), IU11303 (*S. mitis*), IU13333 (*E. faecalis*), for full genotypes see *SI Appendix, Table S1*.

### SI APPENDIX, SUPPLEMENTAL FIGURES AND LEGENDS

**Fig. S1.** Schematic model of PG synthesis in *Spn* by two PG synthesis machines that separate during cell division. The septal machine contains bPBP2x:FtsW and other proteins (pink ovals), which synthesize cross-wall PG that divides the daughter cells. The peripheral machine contains bPBP2b:RodA (green ovals), PG remodeling hydrolases (yellow ovals), and other proteins, which synthesize lateral PG, elongating daughter cells from the midcell (old cell wall, dark blue; new cell wall, light blue). In pre- and early divisional cells (top), both PG synthesis machines localize to the FtsZ ring (gray dotted lines) at the midcell equator of recently divided cells. During division *(middle)*, both machines remain at midcell. Data in this paper show that the sPG synthesis machine and FtsZ ring locate to the leading edge of the constricting septal annulus, separate from the pPG synthesis machine that form in an outer ring lacking FtsZ. During late division (bottom), both PG synthesis machines converge to a single point to complete cell wall synthesis and daughter cell separation. By this time, FtsZ has left the septum and accumulated in rings at the equators of the daughter cells. The sPG and pPG synthesis machine proteins are recruited to these equatorial rings which become the midcell septa in the next round of cell division.

**Fig. S2.** Localization of *(A)* isfGFP-bPBP2x, (*B*) sfGFP-bPBP2b, (*C*) iHT-bPBP2x, (*D*) iHT-bPBP2b, (*E*) iHT-bPBP2x (C+Y), (*F*) native bPBP2x, (*G*) FtsX′-isfGFP-FtsX′, and (*H*) FtsZ-sfGFP in vertically oriented *Spn* cells labeled with an FDAA. Cells were grown in BHI broth, except for (*E*), labeled as indicated below, fixed, and prepared for imaging of vertical cells by 3D-SIM as described above in *SI Appendix*, *Experimental Procedures.* (*A) and (B*) Additional representative images of cells from the experiments described in Fig. 2A. Each row is a different cell (not a time course) that is arranged retroactively by division stage going from early (top) to later (bottom). Cells expressing (*A*) isfGFP-bPBP2x (IU11157; cyan) or (*B*) sfGFP-bPBP2b (IU9965; cyan) were labeled with 125 μM TADA (red) for 2.5 min. (*C*), (*D*), and (*E*). Cells expressing (C) iHT-bPBP2x (IU14927) (top row, merged images of separate cells; bottom row, merged and separate images of a single cell), (D) iHT-bPBP2b (IU16553) (merged images of different cells), or (*E*) iHT-bPBP2x (IU14927) (each row, merged and separate images of two cells grown in C+Y medium instead of BHI broth) were labeled with 500 nM of the HT ligand JF549 (red) for 30 min, washed, and labeled with 400 μM HADA (cyan) for 2.5 min. Cellular amounts of tagged proteins produced relative to the untagged WT protein (see Fig. S3C) are in parentheses. (*F*) WT cells (IU1945) or a Δ*pbp1b* mutant (E193) expressing untagged WT bPBP2x labeled with the β-lactone 7FL probe (green) specific for bPBP2x, viewed by 3D-SIM in horizontally oriented cells at mid- to late-division stages. Two cells of each strain are shown. 7FL predominantly forms a covalent bond with *Spn* bPBP2x and to a lesser extent with aPBP1b (7). Δ*pbp1b* mutants do not show growth or morphology phenotypes under these culture conditions. (*G)* Cells expressing FtsX′-isfGFP-FtsX′ (IU16121; cyan) labeled with 125 μM TADA (red) for 2.5 min. Each row shows representative images of merged and separate cells at different stages of division. The white arrowhead indicates an equatorial ring that is not part of the mature septal ring, but was acquired in the same imaging plane due to a partially vertical orientation. Scale bars = 1 μm. (*H*) Cells expressing FtsZ-sfGFP (IU9985; green) were labeled with 125 μM TADA (red). Scale bars = 1 μm.

**Fig. S3.** Cells expressing sfGFP or HT fusion proteins show minimal growth and morphology defects, and fusion proteins remain intact in cells. The following strains were analyzed: WT (IU1824); isfGFP-bPBP2x (IU11157); iHT-bPBP2x (IU14927); sfGFP-bPBP2b (IU9965); iHT-bPBP2b (IU15928); iHT-bPBP2b//$P_{Zn}$-iHT-bPBP2b (IU16553); FtsX′-isfGFP-FtsX′ (IU16121). Full strain genotypes are in *SI Appendix, Table S1*. (*A*) Growth curves in BHI broth of strains expressing fusion proteins of bPBP2x (left), iHT-bPBP2b (middle), and sfGFP-bPBP2b and FtsX′-isfGFP-FtsX′ (right). Strains in the middle panel were grown in BHI broth supplemented with 0.3 mM $ZnCl_2$ and 0.03 mM $MnSO_4$ (filled symbols) to induce the expression of iHT-bPBP2b in strain (IU16553). For each strain, a representative growth curve from three or more independent biological replicates is shown. Mean growth rates and growth yields (± SD) are shown below the graphs. (*B*) 2D-epifluorescece and phase-contrast microscopy was performed on cells expressing fusion proteins of bPBP2x (left), iHT-bPBP2b (middle), and sfGFP-bPBP2b and FtsX′-isfGFP-FtsX′ (right), grouped underneath and colored to correspond with the growth curves in (*A*). All cells were labeled, fixed and prepared for vertical cell imaging as described for 3D-SIM, but were imaged using 2D-epifluorescence and phase-contrast microscopy (see *SI Appendix*, *Experimental Procedures*). For each strain, a representative field is shown. (*C*) Quantitative western blots of cell lysates from strains expressing fusion proteins of bPBP2x (*top left*), FtsX (top right), and bPBP2b (bottom) were performed using native antibodies as described in *SI Appendix*, *Experimental Procedures*. Descriptions of samples are found at the tops of each lane. +, untagged protein; iHT, isfGFP, and sfGFP, fusion proteins; Δ, Δ*pbp2b* negative control strain (IU9783, *SI Appendix, Table S1*). Along the sides of blots, arrows point to expected molecular masses of each fusion protein, as well as untagged proteins of interest. Within the blot, the amount of sample loaded (total µg of protein extract) is shown in blue. The protein of interest is quantified at the bottom of each lane, relative to the untagged protein (defined as 1; ≡1). Large numbers indicate the mean relative protein amount ± SD from two or more independent biological replicates for a particular fusion protein. Standard curves (to the right of blots) were constructed by loading various amounts of lysate from the same sample and plotting the chemiluminescent signal detected.

**Fig. S4.** Distributions of ring diameters (*A*) and number of nodes per ring (*B*) of mutant strains. Measurements were performed using a custom Matlab VIMA-GUI to analyze FDAA nodes within rings (see *SI Appendix*, *Experimental Procedures*). Graphs show median (interquartile range; whiskers, $5^{th}$-$95^{th}$ percentile; +, mean). Means ± SD are compiled in Table S3. Differences in means relative to WT were determined by one-way ANOVA with Bonferonni's multiple comparison posttest (GraphPad Prism). **, $P<0.01$; ***, $P<0.001$.

**Fig. S5.** Node amount, but not spatial overlap, is positively correlated between TADA nodes and either sfGFP-bPBP2b or Boc-FL nodes in predivisional *Spn* cells. A custom Matlab VIMA-GUI was used to analyze the number and spatial overlap of nodes in predivisional rings from cells expressing sfGFP-bPBP2b (IU9965) and labeled with 45.5

μM of TADA for 17 s (*A* and *B*), or WT cells (IU1945) labeled with TADA and then with 2 μg/mL of Boc-FL for 21 s (*C* and *D*). After labeling, cells were fixed and prepared for imaging of vertical cells using 3D-SIM as described in *SI Appendix*, *Experimental Procedures*. (*A* and *C*) For each measured cell, the number of TADA nodes was plotted against the number of sfGFP-bPBP2b (*A*, *n*=48 cells) or Boc-FL nodes (*C*, *n*=18 cells). (*B* and *D*) For each measured cell, the spatial correlation (the amount of overlap between different types of nodes) was determined for TADA nodes versus sfGFP-bPBP2b (*B*, *n*=48 cells) or Boc-FL nodes (*D*, *n*=18 cells). A coefficient value of -1 indicates a perfect negative correlation, a value of 1 indicates a perfect positive correlation, and a value of 0 indicates no correlation. Data shown are from two independent biological replicates.

**Fig. S6.** Colocalization of cephalosporin conjugated to TAMRA (Ceph-CT) and fluorescent-bocillin (Boc-FL) in horizontally oriented, predivisional *Spn* cells. WT cells (IU1945) were labeled with Ceph-CT (red), then Boc-FL (green), followed by washing and fixation as described in *SI Appendix, Additional Data*. Over 60 cells from more than three independent biological replicates were imaged, and a representative image is shown. Lateral images in the top row are rotated 90° to produce the images in the bottom row.

**Fig. S7.** Tandem pulse labeling with three different colored FDAAs results in regular, displaced nodal patterns for each color of FDAA probe. WT cells (IU1945) were briefly and sequentially labeled with the three indicated FDAAs, fixed, and prepared for imaging of vertical cells using 3D-SIM as described in *SI Appendix*, *Experimental Procedures*. White arrows highlight displacement of nodal labeling for each different colored FDAA. Each row shows a representative cell from two independent biological replicates (*n* = 10 total cells analyzed).

### SI APPENDIX, REFERENCES

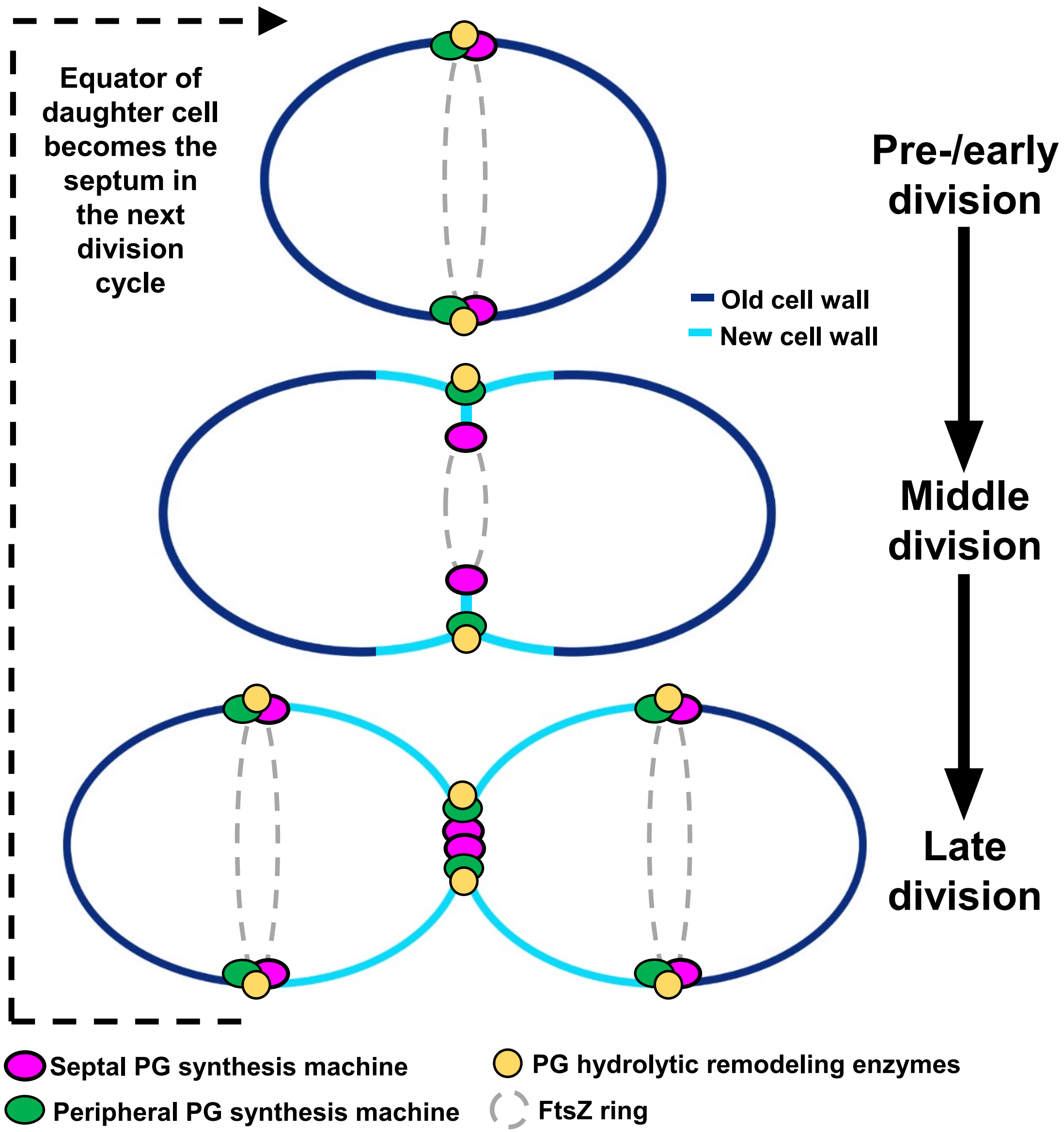
Equator of daughter cell becomes the septum in the next division cycle
Pre-/early division
Middle division
Late division
Old cell wall
New cell wall
Septal PG synthesis machine
Peripheral PG synthesis machine
PG hydrolytic remodeling enzymes
FtsZ ring

Fig. S1

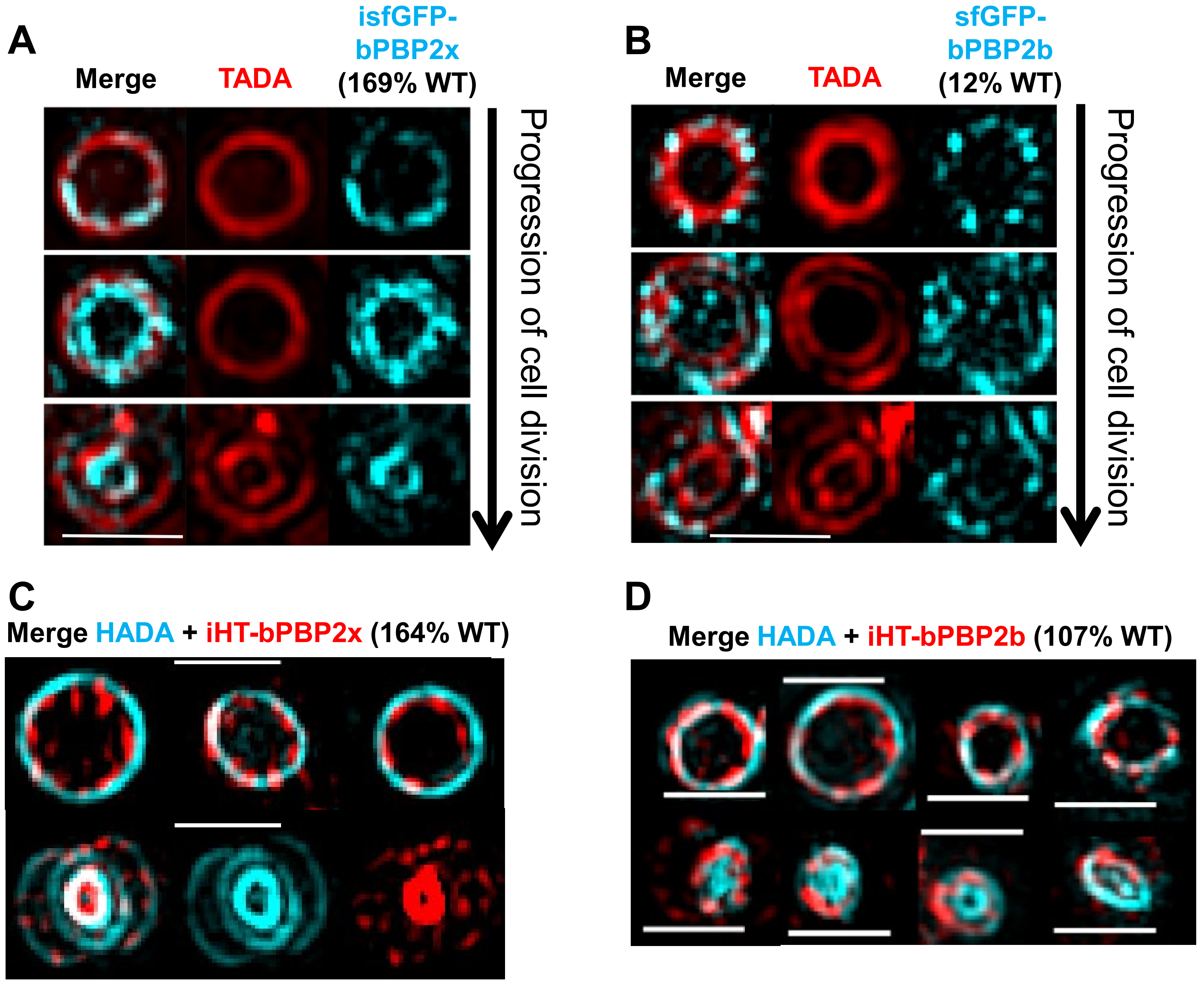


**Fig. S2 (continued on next pages)**

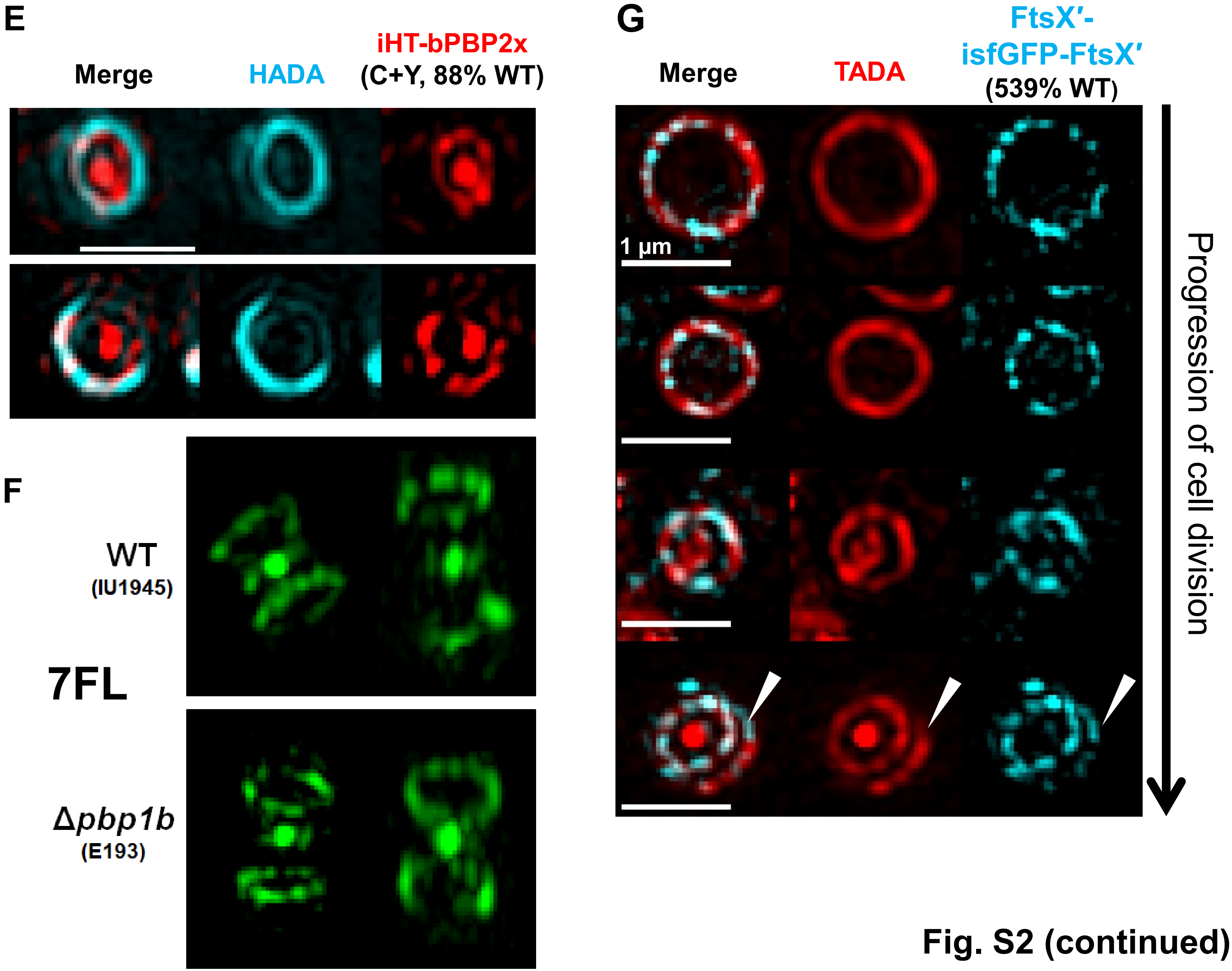


Fig. S2 (continued)

H

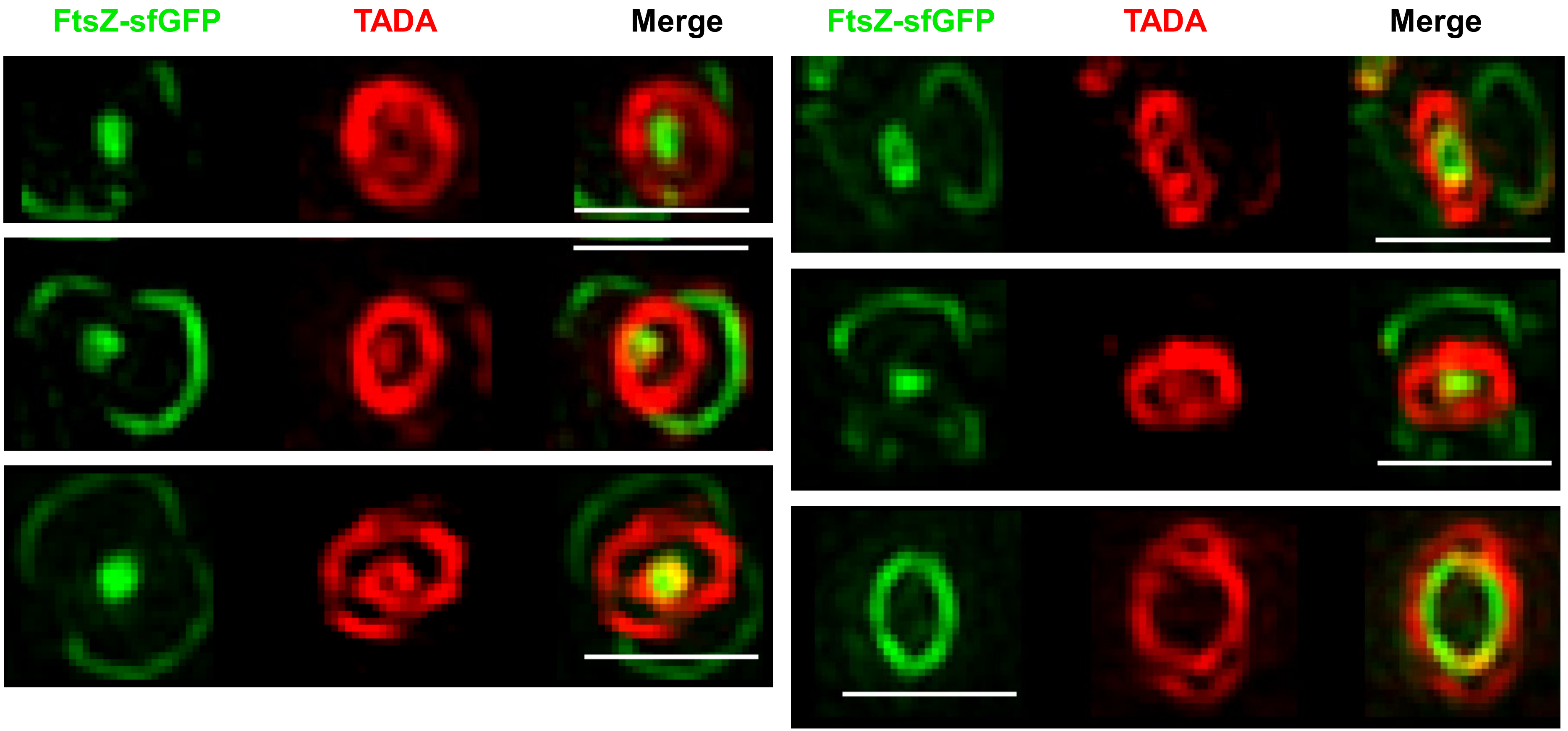


Fig. S2 (continued)

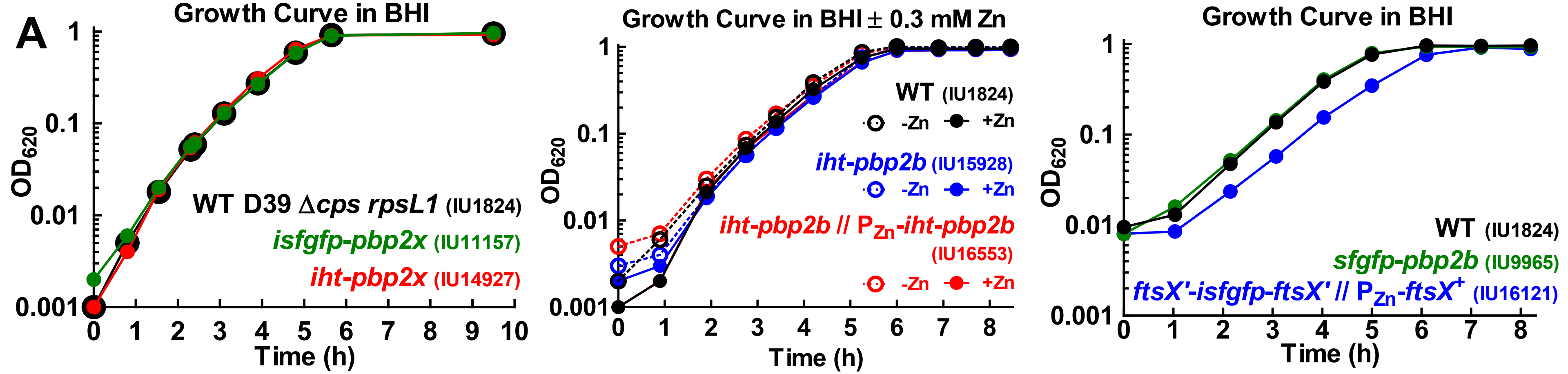


| *n* = 4 | Mean Doubling Time (min ± SD) | Mean Final Yield (OD ± SD) |
|---|---|---|
| WT | 40.3 ± 3.3 | 0.978 ± 0.034 |
| *igfp-2x* | 40.6 ± 5.4 | 0.969 ± 0.051 |
| *iht-2x* | 39.2 ± 1.4 | 0.962 ± 0.034 |

| *n* = 3 | | Mean Doubling Time (min ± SD) | Mean Final Yield (OD ± SD) |
|---|---|---|---|
| WT | -Zn | 40.2 ± 3.8 | 0.993 ± 0.007 |
| | +Zn | 41.5 ± 3.4 | 0.943 ± 0.038 |
| *iht-2b* | -Zn | 39.1 ± 3.4 | 0.97 ± 0.027 |
| | +Zn | 41.4 ± 3.4 | 0.919 ± 0.007 |
| *iht-2b// iht-2b* | -Zn | 44.2 ± 4.9 | 0.982 ± 0.026 |
| | +Zn | 42.8 ± 3.3 | 0.95 ± 0.037 |

| *n* = 4 | Mean Doubling Time (min ± SD) | Mean Final Yield (OD ± SD) |
|---|---|---|
| WT | 38 ± 4.6 | 0.994 ± 0.025 |
| *gfp-2b* | 36.9 ± 4.2 | 0.959 ± 0.029 |
| *ftsX'-gfp-X'* | 38.6 ± 2.3 | 0.92 ± 0.029 |

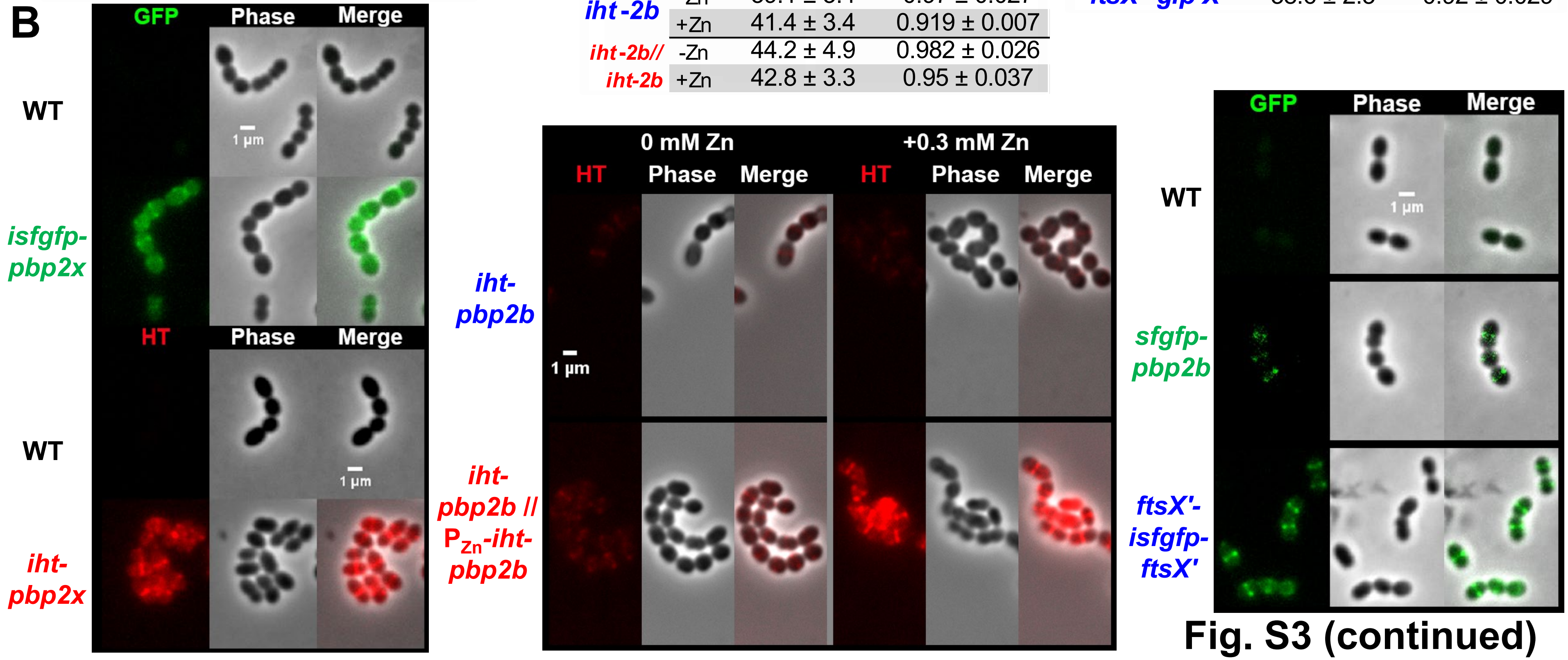


**Fig. S3 (continued)**

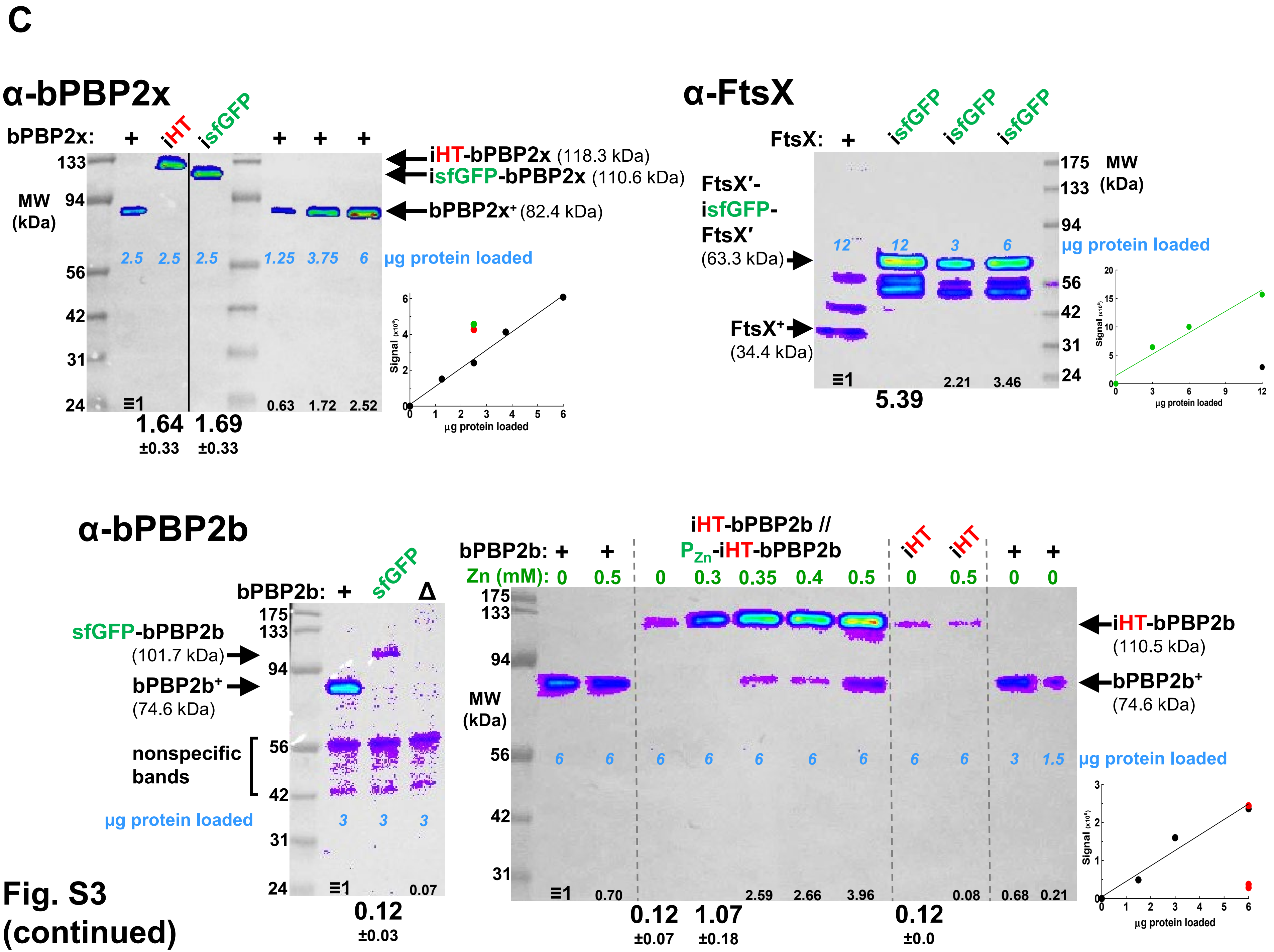


Fig. S3 (continued)

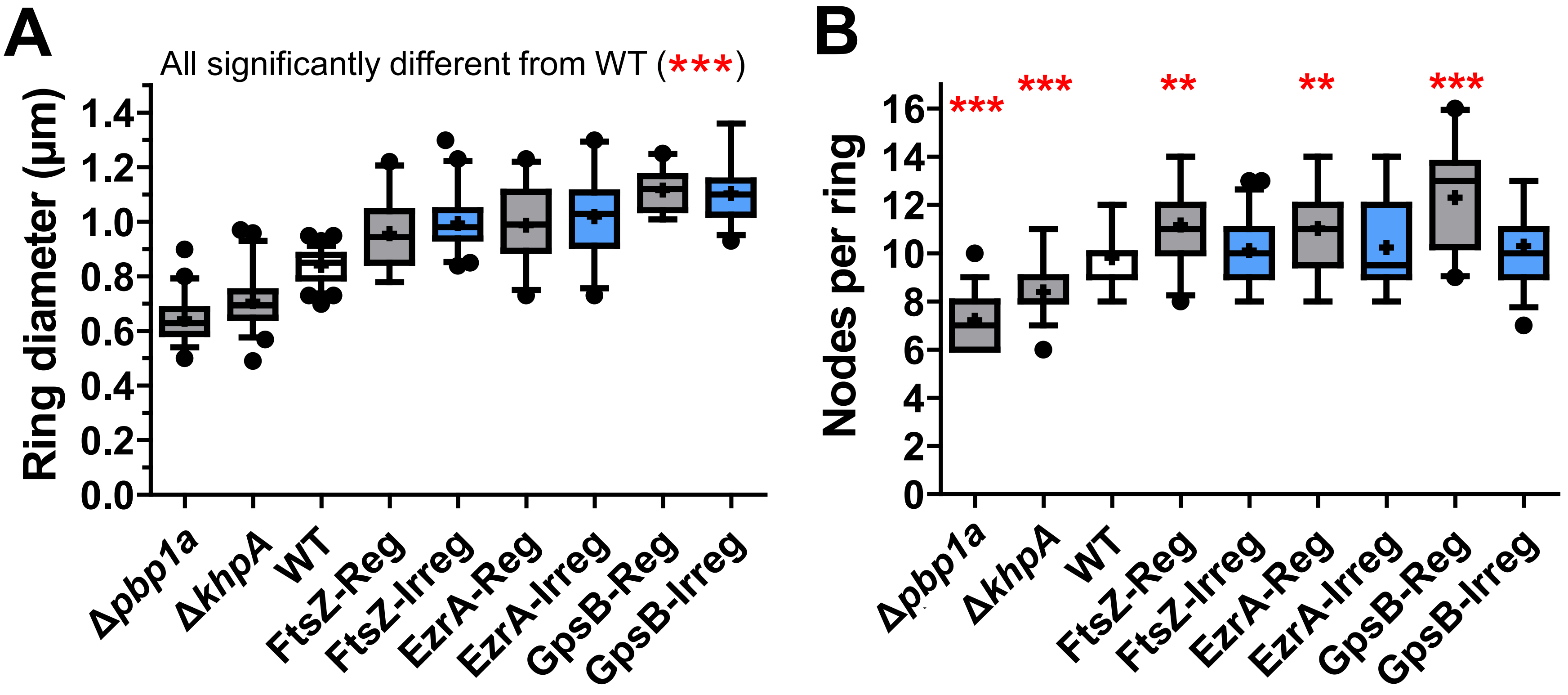
A
All significantly different from WT (***)
Ring diameter (µm)
1.4
1.2
1.0
0.8
0.6
0.4
0.2
0.0
Δpbp1a
ΔkhpA
WT
FtsZ-Reg
FtsZ-Irreg
EzrA-Reg
EzrA-Irreg
GpsB-Reg
GpsB-Irreg
B
Nodes per ring
***
***
**
**
***
16
14
12
10
8
6
4
2
0
Δpbp1a
ΔkhpA
WT
FtsZ-Reg
FtsZ-Irreg
EzrA-Reg
EzrA-Irreg
GpsB-Reg
GpsB-Irreg

Fig. S4

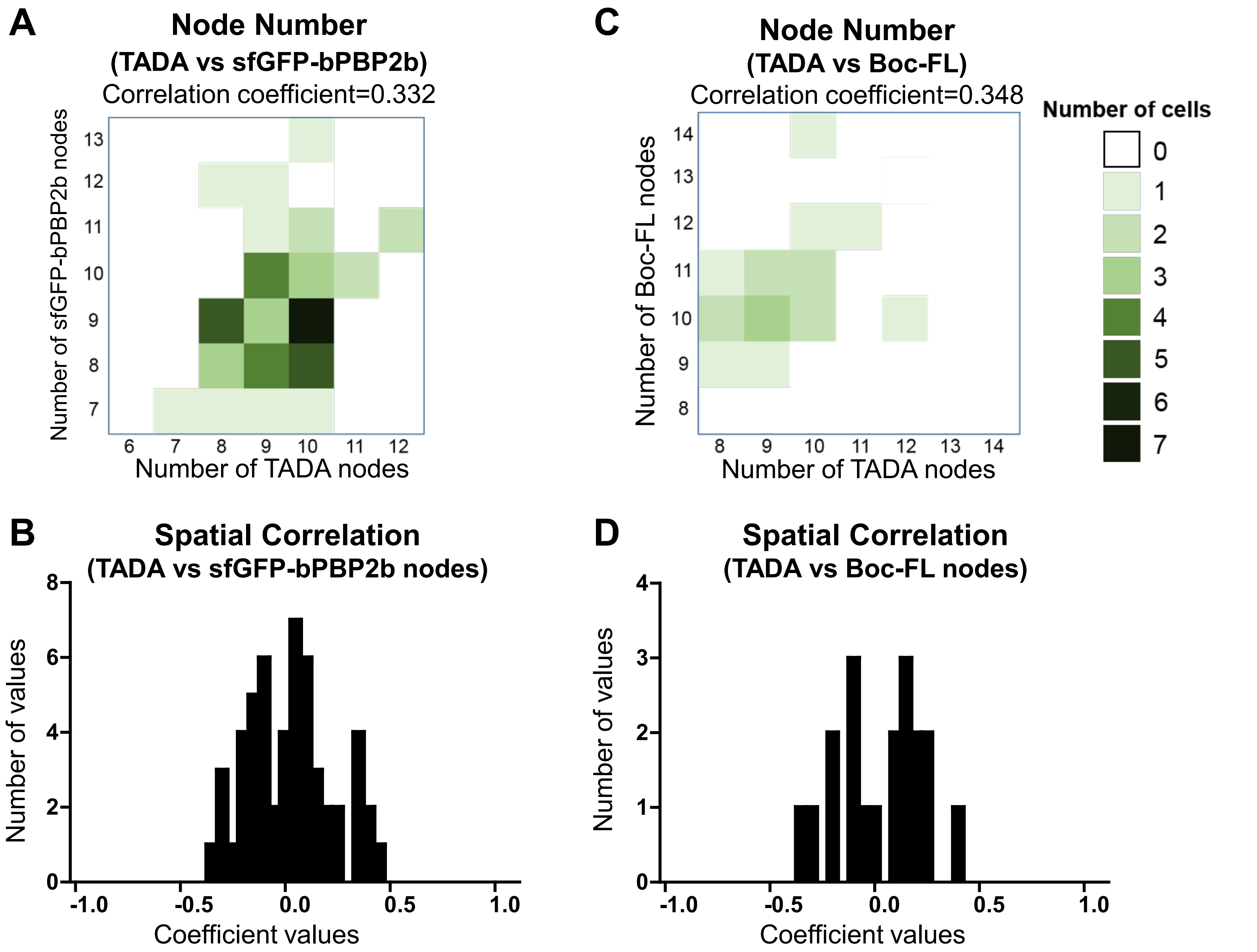
A
Node Number
(TADA vs sfGFP-bPBP2b)
Correlation coefficient=0.332
Number of sfGFP-bPBP2b nodes
Number of TADA nodes
C
Node Number
(TADA vs Boc-FL)
Correlation coefficient=0.348
Number of Boc-FL nodes
Number of TADA nodes
Number of cells
B
Spatial Correlation
(TADA vs sfGFP-bPBP2b nodes)
Number of values
Coefficient values
D
Spatial Correlation
(TADA vs Boc-FL nodes)
Number of values
Coefficient values

**Fig. S5**

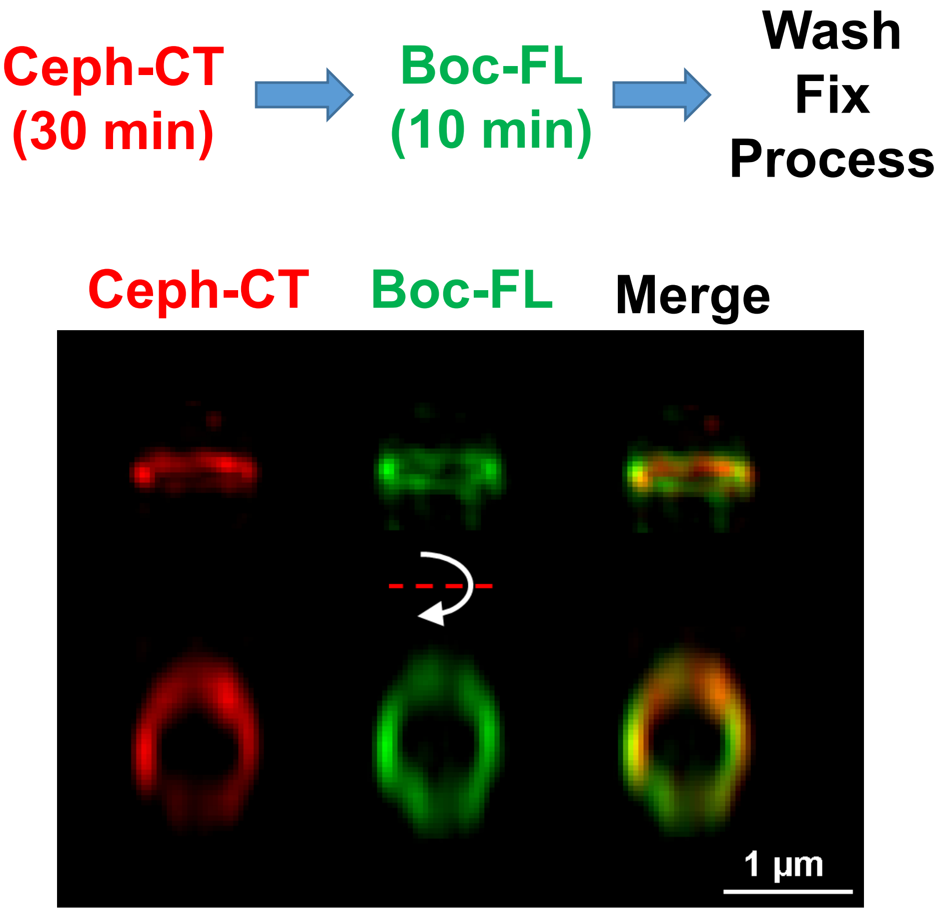
Ceph-CT
(30 min)
Boc-FL
(10 min)
Wash
Fix
Process
Ceph-CT
Boc-FL
Merge
1 µm

Fig. S6

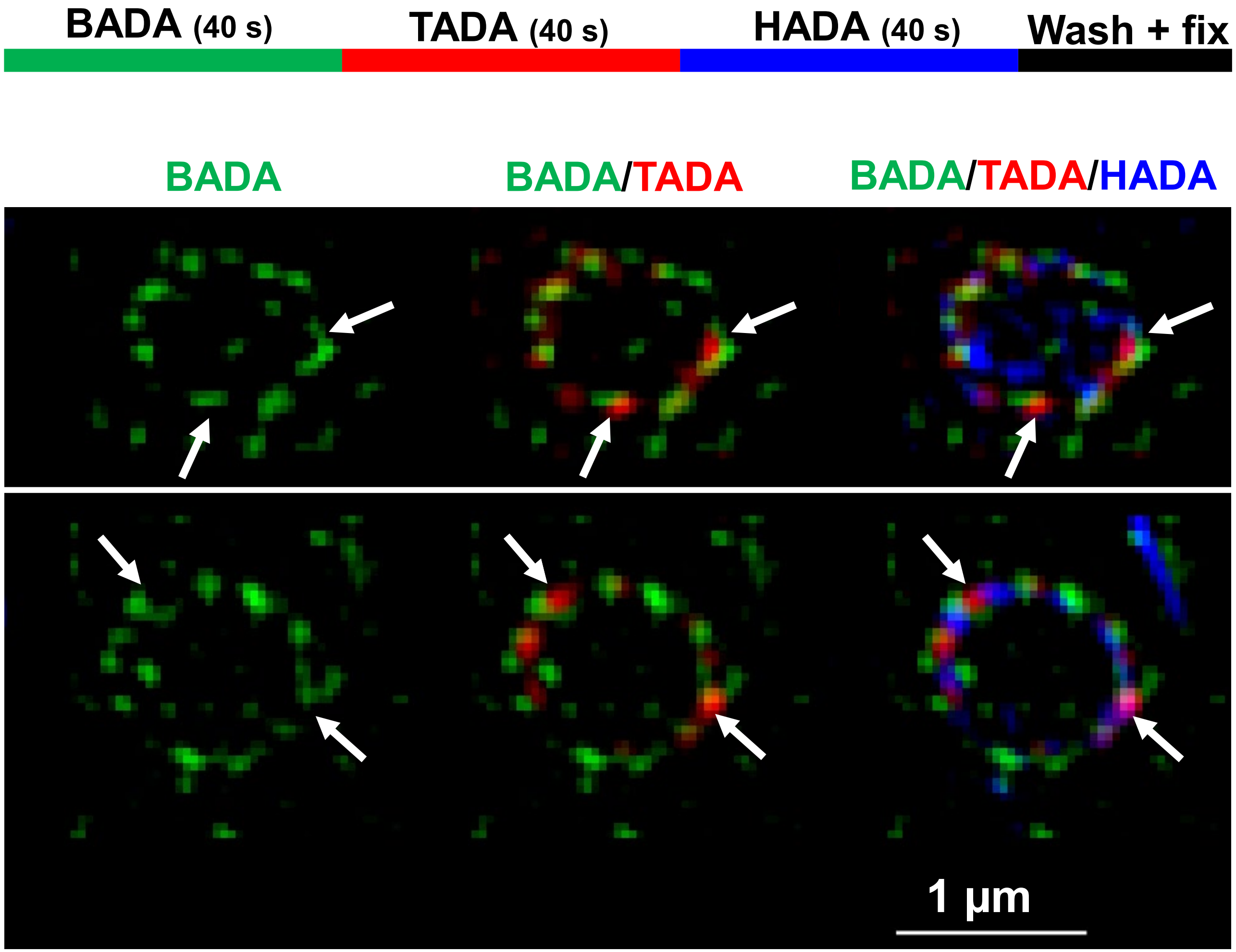

BADA (40 s)
TADA (40 s)
HADA (40 s)
Wash + fix
BADA
BADA/TADA
BADA/TADA/HADA
1 µm


Fig. S7
